# Auxin Plays a Role in the Adaptation of Rice to Anaerobic Germination and Seedling Establishment

**DOI:** 10.1101/2022.05.05.490789

**Authors:** Kuo-Wei Lee, Jeremy J.W. Chen, Chung-Shen Wu, Ho-Chun Chang, Hong-Yue Chen, Hsin-Hao Kuo, Ya-Shan Lee, Yan-Lun Chang, Hung-Chia Chang, Shiau-Yu Shiuea, Yi-Chen Wu, Yi-Cheng Ho, Peng-Wen Chen

## Abstract

Auxin is well known to stimulate coleoptile elongation and rapid seedling growth in the air. However, its role in regulating rice germination and seedling establishment under submergence is largely unknown. Previous studies have shown that excessive levels of IAA frequently cause the inhibition of plant growth and development. In the present study, the high-level accumulation of endogenous IAA is observed under submergence in the dark, stimulating rice coleoptile elongation but limiting the root and primary leaf growth during anaerobic germination (AG). We found that oxygen and light can reduce IAA levels, promote the seedling establishment and enhance rice AG tolerance. miRNA microarray profiling and RNA gel blot analysis results show that the expression of miR167 is negatively regulated by submergence; it subsequently modulates the accumulation of free IAA through the miR167-ARF-GH3 pathway. The *OsGH3-8* encodes an IAA-amido synthetase that functions to prevent free IAA accumulation. Reduced miR167 levels or overexpressing *OsGH3-8* increase auxin metabolism, reduce endogenous levels of free IAA and enhance rice AG tolerance. The present study reveals that poor seed germination and seedling growth inhibition resulting from excessive IAA accumulation would cause intolerance to submergence in rice, suggesting that a certain threshold level of auxin is essential for rice AG tolerance.

## Introduction

The conventional puddled rice cultivation system has become increasingly unsustainable due to global climate change and socio-economic transition (Sandhu et al., 2021), and direct-seeded rice (DSR) has emerged as a promising alternative to conventional transplantation (Panda et al., 2021). Among major cereals, rice is the only crop that can successfully germinate and grow under submergence. When germinated underwater, rice coleoptile rapidly elongates and reaches a length longer than that of the aerobic coleoptile, whereas root and primary leaf fail to grow. Rapid elongation of hollow coleoptile acts as a snorkel, extending to the water surface, thereby obtaining more O_2_ to support more vigorous seedling establishment. As a result, strong and healthy seedlings exhibit greater tolerance to submergence during germination and early seedling growth (Magneschi and Perata, 2009). Conversely, poor germination and seedling establishment of intolerant rice varieties may affect the acquisition of sufficient O_2_ and limit energy supply, leading to the inability to survive in submerged fields (Hsu and Tung, 2015). Thus, anaerobic germination (AG) tolerance and the establishment of vigorous seedlings under submergence are crucial traits for developing rice varieties, which can be applied in the large-scale adoption of a direct seeding system (Kretzschmar et al., 2015).

In rice, sugar availability has been considered one of the critical factors for tolerance to submergence (Loreti et al., 2003b). Rice seeds can germinate and produce α-amylase enzymes required for starch degradation even without oxygen. Other cereal grains fail to mobilize the starchy endosperm anaerobically and, therefore, cannot germinate and grow underwater (Perata, 1993; Guglielminetti et al., 1995). Under flooded conditions, the oxygen supply is limited and aerobic respiration is reduced, leading to an increase in glycolytic flux and fermentative metabolism (so-called Pasteur effect) and faster consumption of the available carbohydrates. Consequently, rapid depletion of soluble carbohydrates and energy deprivation occur under submergence. Previous studies have shown that sucrose non-fermenting 1 (Snf1)-related protein kinase 1 (SnRK1A), the plant global energy and stress sensor, plays a pivotal role in response to sugar and oxygen deficiency in rice (Lu et al., 2007; Lee et al., 2009). Submergence triggers sugar starvation and induces mRNA accumulation of *calcineurin B-like* (*CBL*) *protein-interacting protein kinase 15* (*CIPK15*), thereby enhancing the accumulation of SnRK1A proteins. These two proteins interact and induce the MYBS1 transcription factor, subsequently activating the expression of starvation-induced α-amylase gene, *αAmy3*/*RAmy3D* (Lee et al., 2009). The CPK15-SnRK1A-MYBS1-α-amylase signaling pathway has been demonstrated to be critical to sugar and oxygen deficiency in rice (Lee et al., 2009; Lee et al., 2014; Wu et al., 2014b). A trehalose-6-phosphate phosphatase gene, *OsTPP7*, has been identified and demonstrated to promote the seedling establishment and enhance the AG tolerance in rice (Kretzschmar et al., 2015). *OsTPP7* is involved in trehalose-6-phosphate (T6P) metabolism and catalyzes the conversion of T6P to trehalose. Reduced T6P level prevents the inhibition of SnRK1A activity and allows for increased starch mobilization in the form of readily fermentable sugar, ultimately enhancing coleoptile elongation and embryo germination (Kretzschmar et al., 2015).

Phytohormones play a crucial role in regulating growth and developmental process and signaling networks involved in plant responses to environmental stresses, including flooding (Jackson, 2008; Khan et al., 2012). Ethylene is the primary regulator of plant adaptation to flooding. It has been previously demonstrated that ethylene is independent of the elongation of rice coleoptile growing in the complete absence of oxygen because oxygen is required for the generation of ethylene (Pearce and Jackson, 1991; Pearce et al., 1992). Gibberellin is considered essential in regulating the expression of α-amylase genes, which catalyze hydrolytic starch degradation during cereal seed germination in the air. However, under anoxic conditions, starch degradation through the gibberellin-induced α-amylase pathway fails to function properly because oxygen is also required for gibberellin biosynthesis, and rice became gibberellin insensitive under anoxic or hypoxic conditions (Loreti et al., 2003a). Auxin is the most abundant plant hormone that plays an essential role in diverse aspects of plant growth and development, including cell division, elongation, differentiation, organ morphogenesis, gravitropic and phototrophic responses, and environmental changes (Leyser, 2018; Gallei et al., 2020). Auxin is well known for promoting coleoptile elongation and rapid seedling growth during germination (Gallei et al., 2020), but little is known about its role in rice germination and seedling establishment under submergence. A recent study has shown that higher auxin availability and transport are involved in stimulating the coleoptile elongation in *japonica* rice under submergence in the dark. Their results also showed that auxin biosynthesis and auxin influx carrier AUX1 regulated the final length of rice coleoptile underwater (Nghi et al., 2021). In higher plants, excessive auxin accumulation frequently causes the inhibition of growth and development, such as root growth inhibition (Yin et al., 2011; Fendrych et al., 2018); thus, the optimal level of endogenous auxin, particularly IAA, must be tightly controlled. Plants use several mechanisms to maintain IAA homeostasis, including biosynthesis, degradation, transport, and conjugate formation. Previous reports have shown that IAA conjugates, such as IAA-Ala, IAA-Leu, and IAA-Phe, serve as storage conjugates and could be hydrolyzed back to free IAA via amino acid conjugate hydrolases. IAA-Asp and IAA-Glu are thought to be precursors for the degradation pathway, and oxIAA is one of the major degradation products of IAA (Ludwig-Müller, 2011). In addition, light is critical for plant survival and plays a role in regulating the auxin homeostasis. Previous studies have demonstrated that the growth of seedlings under high R: FR (the ratio between red and far-red light) conditions or high photosynthetically active radiation (PAR) are mediated through phytochrome B (phyB) to reduce IAA levels and inhibit auxin response when compared to those in darkness or low R: FR conditions (Zhang et al., 2014).

MicroRNA-mediated signaling has been shown to be involved in plant growth and development and stress responses through the auxin signaling pathway (Meng et al., 2010; Sanan-Mishra et al., 2013); however, the role of miRNAs in rice’s response to AG tolerance is still largely unknown. A rice microRNA, miR393, has been demonstrated to be involved in coleoptile elongation under submergence in the dark (Guo et al., 2016). miR393a was found to negatively regulate the auxin signaling pathway by post-transcriptional cleavage of *transport inhibitor response1* (*TIR1*) mRNA, an auxin receptor (Bian et al., 2012). The expression of miR393a is repressed by submergence, and the auxin response is induced, which in turn stimulates coleoptile elongation. Overexpression of miR393a inhibits stomatal development, reduces the length of coleoptile, and significantly increases the free IAA level in rice coleoptile. It was suggested that miR393a may be involved in maintaining auxin homeostasis and auxin signaling to modulate seed germination and early seedling growth under submergence (Guo et al., 2016). In addition to miR393a, miR167 has been shown to play a role in plant growth and development process through auxin signaling (Meng et al., 2010; Sanan-Mishra et al., 2013). miR167 was shown to negatively regulate four *auxin response factor* (*ARF*) genes in rice, *OsARF6, OsARF12, OsARF17, and OsARF25* (Liu et al., 2012), and regulate intracellular free IAA levels through the miR167-ARF-GH3 pathway (Yang et al., 2006). MiR167-directed *OsARFs* mRNA cleavage is accompanied by the down-regulation of *OsGH3* (*glycoside hydrolase 3*) genes expression (Yang et al., 2006). The auxin-responsive *OsGH3* genes, such as *OsGH3-2*, *OsGH3-6*, and *OsGH3-8*, encode an IAA-amido synthetase that functions to prevent excessive accumulation of free IAA and help in maintaining auxin homeostasis (Jain et al., 2006a; Ding et al., 2008; Ludwig-Müller, 2011; Kong et al., 2019). In addition, miR167 also negatively regulated the *IAR3* (*IAA-Ala Resistant3*) gene, which encodes an IAA-amino acid hydrolase and releases bioactive IAA via hydrolysis of amino acid-type IAA conjugates; therefore it was suggested to play a role in regulating the levels of free IAA (Davies et al., 1999; Kinoshita et al., 2012). It was previously demonstrated that miR167 expression was down-regulated by long-term waterlogging in maize roots (Zhai et al., 2013); however, whether miR167-mediated signaling is involved in the regulation of rice germination and seedling establishment under submergence remains elusive.

It is known that the growth and development of rice seedlings are inhibited under submergence compared to those in the air. Root growth inhibition in response to the asymmetrical distribution of auxin to confer positive gravitropism is an essential trait for plants to colonize land and utilize water and nutrients. It has been demonstrated that auxin-mediated root growth inhibition is dependent on intracellular IAA accumulation, and the inhibition was triggered via auxin receptor localized inside the cell (Fendrych et al., 2018). Thus, whether auxin plays a role in regulating rice adaptation to AG and seedling establishment remains to be explored. In the present study, phototropic response studies indicated that rice seedlings are sensitive to endogenous auxin under submergence. We found that auxin is essential for AG and the seedling establishment of rice. Accumulation of high-level endogenous free IAA stimulated rice coleoptile elongation but limited root and primary leaf growth under submergence in the dark. We also found that oxygen and light significantly reduced free IAA levels and enhanced AG tolerance in rice. A submergence-repressible miR167 was identified and it was involved in the metabolic regulation of IAA mediated through the miR167a-ARF-GH3 pathway. Our results demonstrate that reduced IAA levels promote the seedling establishment and enhance rice AG tolerance, suggesting that a certain threshold level of endogenous auxin is required for rice germination and early seedling growth under submergence.

## Results

### Oxygen and light reduce the accumulation of free IAA and enhance the seedling establishment during AG

To examine the effect of oxygen on the morphological traits of seedlings and the accumulation of endogenous free IAA during AG, rice seeds (*Oryza sativa* L. cv. Tainung 67) were germinated and grown in the dark for 6 days in the air or under submergence in the dark as described in *Materials and methods*, and the concentration of free IAA and its conjugates in the aerobic and hypoxic seedlings were quantified using UPLC-ESI-MS/MS analysis. Seeds germinated in aerobic environment grew root, primary leaf, and coleoptile, whereas seeds germinated in stagnant water had elongated and overgrown coleoptile without root and primary leaf. When oxygen was provided, the morphology of rice seedlings grown under aerated water was similar to that of the aerobic seedlings (Figure 1A). The concentration of free IAA in the hypoxic seedlings grown under stagnant water was 3.9-fold higher than that in the aerobic seedlings; nevertheless, the IAA level only increased slightly in seedlings grown under aerated water. Under stagnant water, the concentration of IAA conjugates, IAA-Ala, in the seedlings was significantly lower, whereas the concentration of IAA-Glu, IAA-Asp, and oxIAA was significantly higher than that of the seedlings grown in the air or under aerated water (Figure 1B). These results were similar to previous findings in rice seedlings and suggested that high-level endogenous free IAA may be responsible for growth inhibition of root and primary leaf in rice seedlings grown under stagnant water, and the metabolism of IAA was shifted to promote the degradation process during AG.

**Figure 1.**
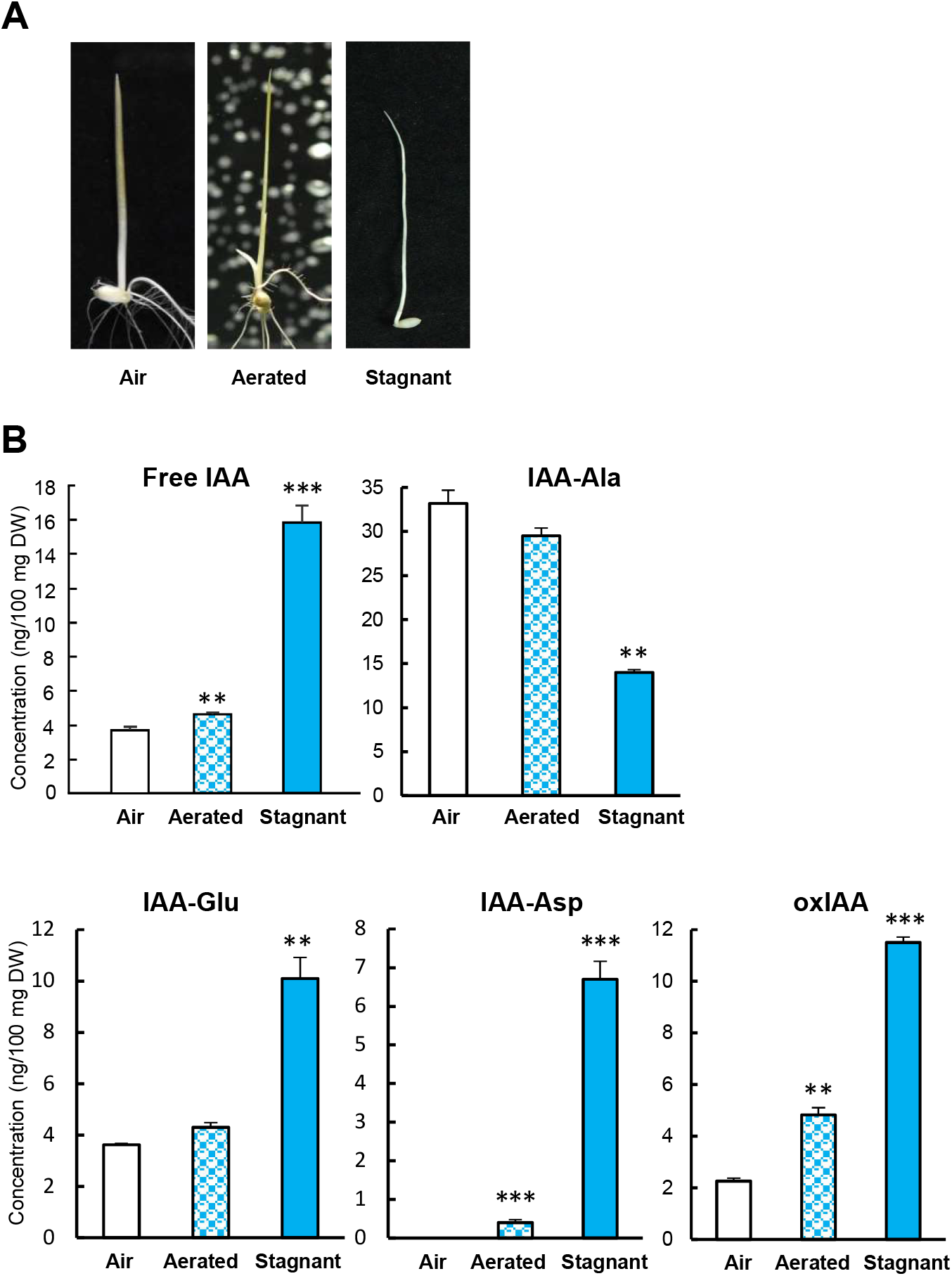
High-level accumulation of endogenous free IAA in hypoxic rice seedlings during anaerobic germination. Rice seeds (*Oryza sativa* L. cv Tainung 67) were germinated and grown under aerobic (Air, *open bar*) conditions or underwater and continuously bubbled with (Aerated, *blue-dotted bar*) or without (Stagnant, *blue bar*) air for 6 days at 28*°*C in the dark. (A) Morphology of 6-day-old rice seedlings. (B) Seedlings were collected and the concentrations of endogenous free IAA, IAA conjugates (IAA-Ala, IAA-Glu, and IAA-Asp) and metabolites (oxIAA) were analyzed by ultra-performance liquid chromatography-electrospray ionization-tandem mass spectrometry (UPLC-ESI-MS/MS). *Error bars* represent the standard error (S.E.) of three replicate experiments for each treatment. Data were analyzed using Student’s t-test, and asterisks indicate significant differences between aerobic and hypoxic rice seedlings (** = *P* < 0.01, *** = *P* < 0.001).

To determine whether light could reduce free IAA level and promote the root and primary leaf growth during AG, rice seeds were germinated and grown in darkness or a light-dark cycle under aerobic or submerged conditions for 7 days, and the level of free IAA and its conjugates were determined. As shown in Figure 2A, light promoted the growth of hypoxic seedlings from only coleoptile elongation to including root and primary leaf growth. In addition, light significantly reduced the level of free IAA in aerobic and hypoxic seedlings, with a 3.3- and 2.3-fold decrease, respectively (Fig. 2B). The level of IAA-Ala was relatively low in the hypoxic seedlings regardless of whether the seedlings were grown in the dark or under light conditions. The level of IAA-Glu, IAA-Asp, and oxIAA in all groups exhibited a positive correlation with the accumulation of free IAA in rice seedlings (Figure 2B). Taken together, these results demonstrate that oxygen and light significantly reduce the endogenous level of free IAA in rice seedlings and greatly enhance the seedling establishment during AG.

**Figure 2.**
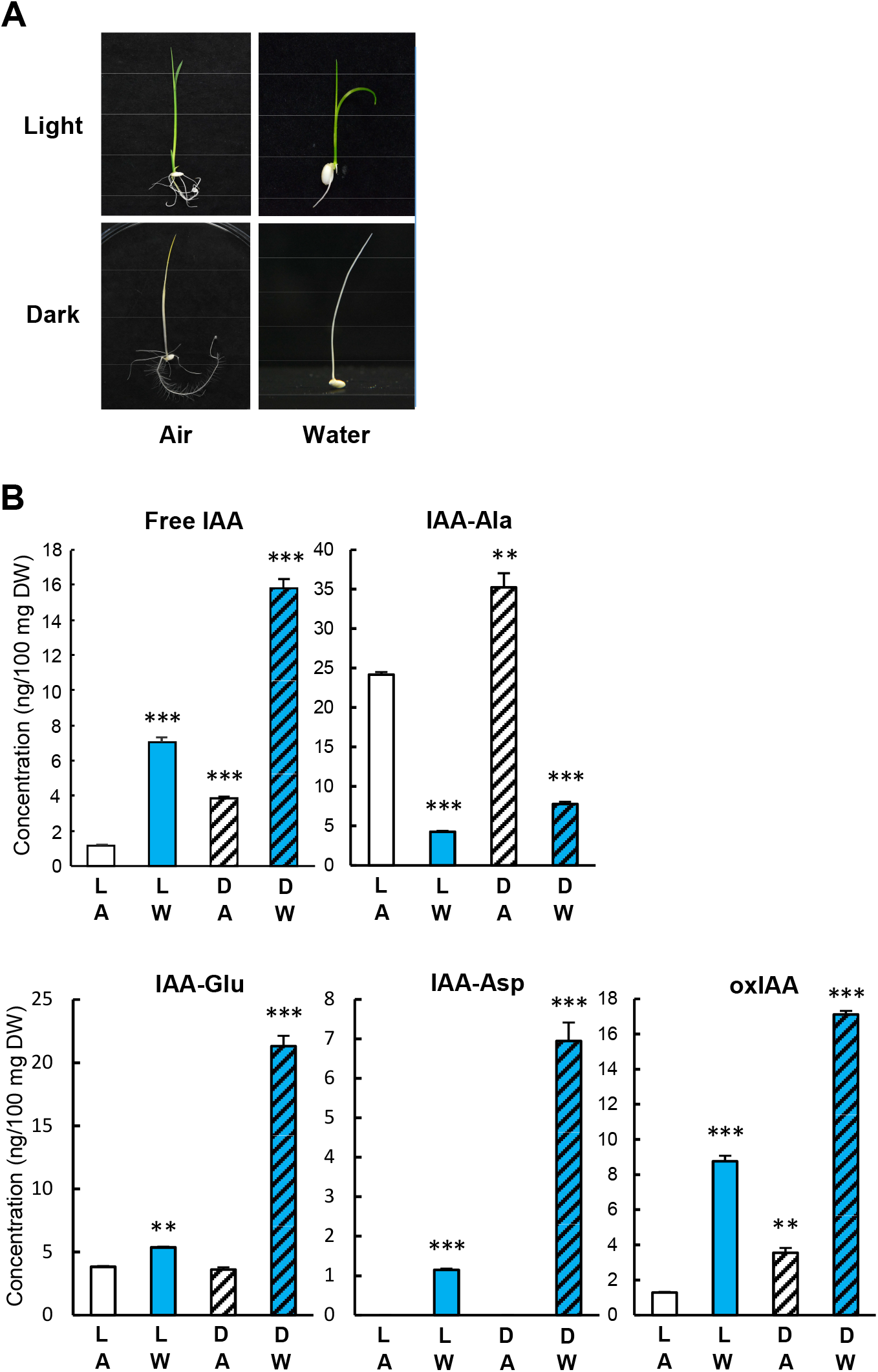
Effect of light on the morphology and accumulation of endogenous free IAA, IAA conjugates, and IAA metabolites in rice seedlings during aerobic and anaerobic germination. Seeds from TNG67 were germinated and grown in a 16-h light (5,000 lux)/8-h dark cycle (Light, L, *open bar*) or in the dark (Dark, D, *hatched bar*) under aerobic (Air, A, *white color*) or submerged (Water, W, *blue color*) conditions at 28*°*C for 7 days. (A) Morphology of 7-day-old rice seedlings grown in aerobic (Air) and anaerobic (Water) condition. (B) Seedlings were collected and the concentrations of endogenous free IAA, IAA conjugates, and IAA metabolites were determined by UPLC-ESI-MS/MS. *Error bars* represent the S.E. of three replicate experiments for each treatment. Student’s t-test was used to analyze the statistical difference between treatment groups and seedlings grown with air and light (L/A). ** = *P* < 0.01, *** = *P* < 0.001.

### The hypoxic seedlings of AG-susceptible variety have higher IAA levels than those of AG-tolerant variety

To determine whether AG intolerance and poor seedling establishment in AG-susceptible rice varieties are due to higher-level accumulation of endogenous free IAA in hypoxic seedlings, seeds from AG-tolerant *japonica* landrace Khao Hlan On (KHO) and Tainung 67 (TNG67) and AG-susceptible *indica* variety IR64 were germinated and grown in a light-dark cycle under aerobic or submerged conditions for 8 days, and the concentrations of free IAA and its conjugates were determined. As shown in Figure 3A, the root length of TNG67 seedlings grown in the air was greater than that of KHO and IR64 at 4, 6, and 8 days after imbibition (DAI), whereas the aerial part showed the opposite result and the aerial part of TNG67 was shorter than that of KHO and IR64 at 4 and 6 DAI. Under submerged conditions, rapid growth and elongation of coleoptile was observed in TNG67 and KHO but not in IR64 at 4 DAI. The hypoxic seedlings of TNG67 grew root and primary leaf at 8 DAI, and the morphology of seedlings was similar to those of KHO, whereas poor seedling establishment was found in AG-susceptible IR64 at 8 DAI. All three rice varieties had higher free IAA levels in hypoxic seedlings than their aerobic counterparts, and the highest IAA level was detected in IR64 (Figure 3B). The highest-level of free IAA was detected at 4 DAI and declined at 6 and 8 DAI in both aerobic and hypoxic seedlings. The concentration of IAA-Ala in hypoxic seedlings of three rice varieties was lower than in aerobic seedlings. On the contrary, the concentrations of IAA-Glu, IAA-Asp, and oxIAA were higher in hypoxic seedlings than in their aerobic counterparts (Figure 3B).

**Figure 3.**
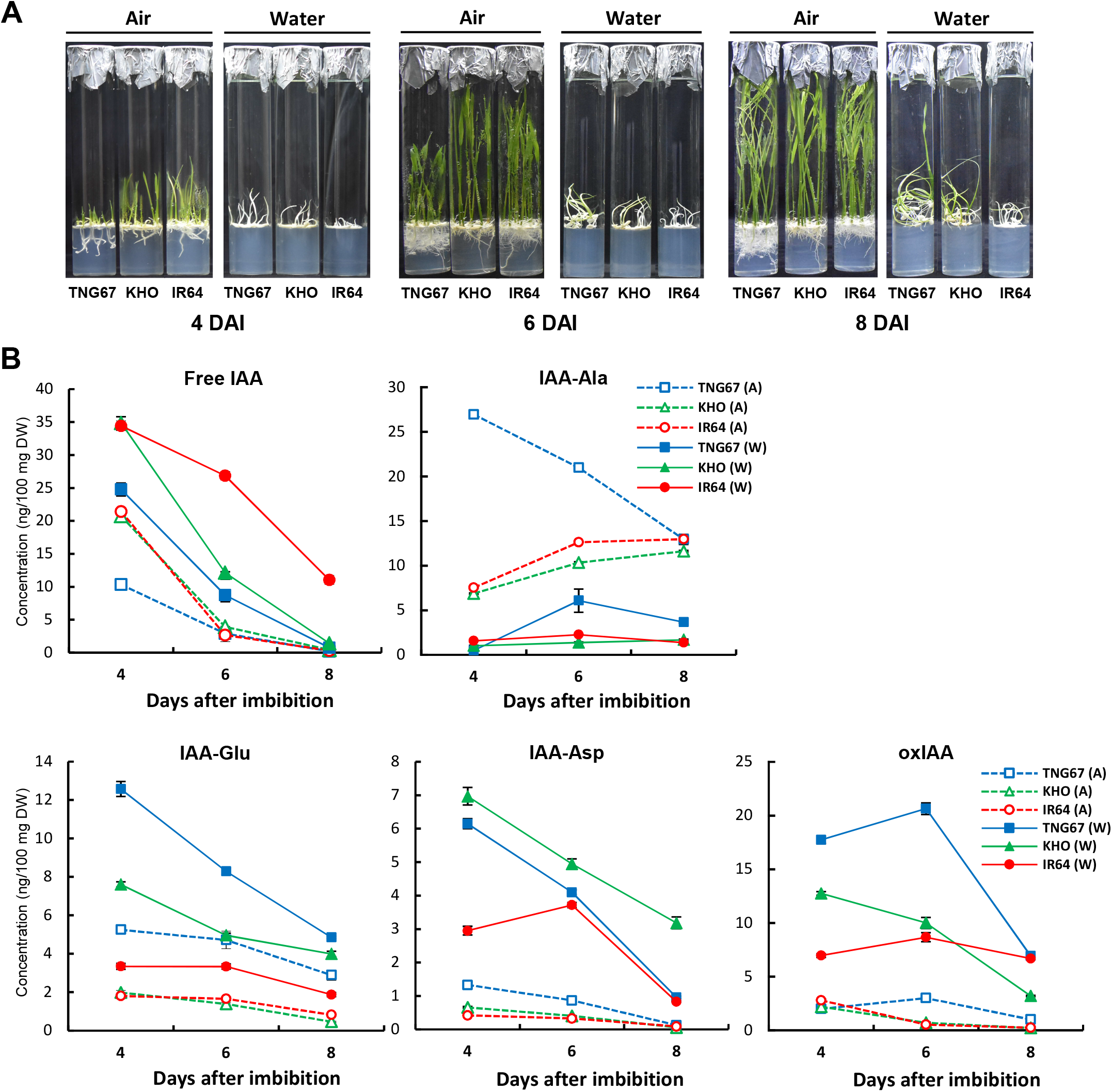
Higher-level accumulation of free IAA is found in hypoxic seedlings of an AG-susceptible variety IR64. Seeds from AG-tolerant *japonica* landrace Khao Hlan On (KHO) and Tainung 67 (TNG67) and AG-susceptible *indica* variety IR64 were germinated and grown in a 16-h light/8-h dark cycle under aerobic (Air) or submerged (Water) conditions for 8 days. (A) Seedling morphology of three rice varieties at 4, 6, and 8 days after imbibition (DAI). (B) Seedlings were collected, and the concentrations of endogenous free IAA, IAA conjugates, and IAA metabolites were determined. *Error bars* represent the S.E. Significance level was analyzed with One-way ANOVA, *P* < 0.01.

### Inhibition of IAA functions enhances AG tolerance in hypoxic seedlings

To evaluate the effect of low oxygen tension on AG tolerance in rice, seeds from TNG67 were germinated and grown under fully embedded conditions for 5 days in a light-dark cycle. Seeds were placed in a test tube and filled with sterile water containing 0 to 2.5 g/L phytagel to mimic reduced oxygen tension. As shown in Supplemental Figure S1, the root, primary leaf, and coleoptile were produced in sterile water containing 0 to 10^-2^ g/L phytagel, and the root length of hypoxic seedlings was increasingly shorter as the phytagel concentration increased. The root and primary leaf failed to grow, and only the coleoptile overgrew when the concentration of phytagel was above 1×10^-2^ g/L. Under such low oxygen tension, coleoptile can elongate up to approximately 3-4 cm. Nevertheless, the coleoptile was unable to turn green, which was essential for the emergence of root and primary leaf. Our result reveals that decreasing oxygen tension underwater negatively affected the rice AG tolerance under fully embedded conditions

To compare the ability of TNG67 and IR64 to resist low oxygen tension, seeds from these two rice varieties were germinated and grown under fully embedded conditions as described above, and filled with sterile water containing 0 to 4 x10^-2^ g/L phytagel. Greening rate, rooting rate, shooting rate, seminal root length, and seedling height were measured daily. In the beginning, the coleoptile of TNG67 elongated and was overgrown. The coleoptile tip turned green at 2-4 DAI and the hypoxic seedlings were rooting and shooting in sterile water containing 0 to 2 x10^-2^ g/L phytagel at 5 DAI (Supplemental Figure S2, A and C). These results also confirm that the greening of the coleoptile plays a pivotal role in the subsequent growth of root and primary leaf. When the concentration of phytagel was above 2×10^-2^ g/L, the coleoptile still overgrew, but the coleoptile failed to turn green and there was no root and primary leaf formation (Supplemental Figure S2, A, B, and C). These results suggested that the rate of greening, rooting, and shooting negatively correlated with increasing concentration of phytagel. At 8 DAI, the seminal root length and seedling height of TNG67 hypoxic seedlings also decreased as the concentration of phytagel increased (Supplemental Figure S2, B and D). The poor seedling establishment of IR64 hypoxic seedlings was shown at 5 and 8 DAI and results were similar to those found with TNG67; all physiological traits of the developmental stage were negatively affected when the concentration of phytagel increased (Supplemental Figure S2, E, F, G, and H).

The above results demonstrated that higher free IAA accumulation in rice hypoxic seedlings leads to poor seedling establishment and reduces the AG tolerance (Figures 1 to 3). To determine whether treatment of free IAA, auxin-action inhibitor, PCIB, and auxin-transport inhibitor, NPA, could affect the AG tolerance, seeds from TNG67 and IR64 were germinated and grown under fully embedded conditions for 10 days in a light-dark cycle. Seeds were placed in a test tube and filled with sterile water containing 2 x 10^-2^ g/L phytagel with or without IAA, PCIB, or NPA. Greening rate, rooting rate, shooting rate, seminal root length, and seedling height were measured daily (Supplemental Figure S3). As shown in Figure 4A, the addition of free IAA significantly reduced the growth of TNG67 hypoxic seedlings in a dose-dependent manner at 5 and 8 DAI. The addition of PCIB, particularly at 20 μM, significantly promoted AG tolerance and enhanced the establishment of solid hypoxic seedlings. The NPA treatment at 0.01 μM also enhanced AG tolerance in TNG67 hypoxic seedlings, but such enhancement was not observed at higher concentrations of NPA, indicating that auxin transport is required for rice germination and seedling growth underwater. A similar result was observed in IR64 hypoxic seedlings and the addition of free IAA inhibited the root and primary leaf formation. The PCIB treatment, particularly at 2.5-10 μM, also enhanced AG tolerance in IR64 hypoxic seedlings (Figure 4B). Unlike TNG67, NPA treatment had no effect on AG tolerance enhancement in IR64 hypoxic seedlings (Fig. 4B). These results indicated that inhibition of IAA functions could promote seedling establishment of TNG67 and IR64, possibly by enhancing their AG tolerance. It also suggests that auxin is essential for both coleoptile elongation and the growth of root and primary leaf and therefore a certain threshold level of endogenous auxin and optimal distribution of auxin are required for rice germination and early seedling growth under submergence.

**Figure 4.**
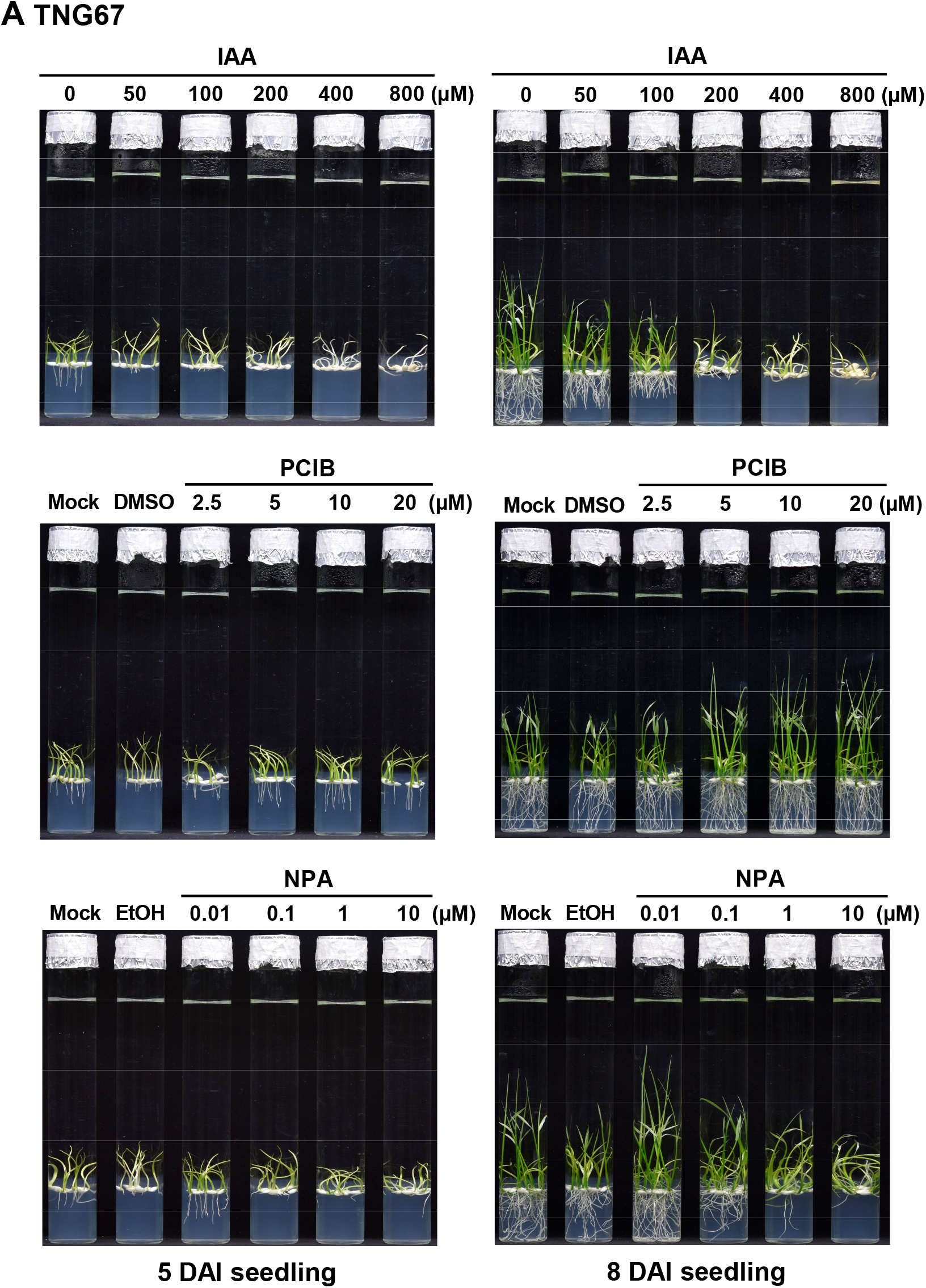
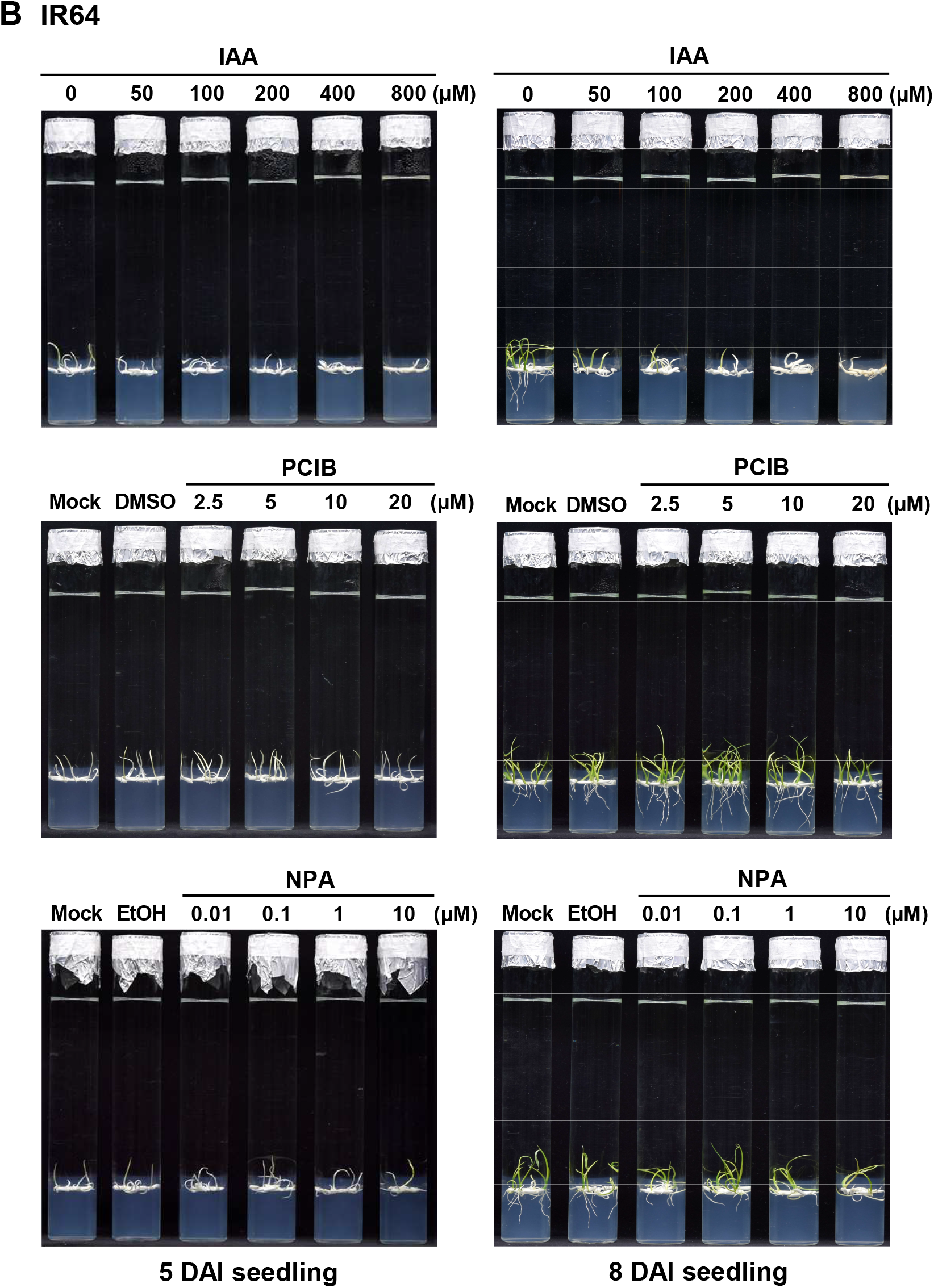
Effect of treatment with different concentrations of free IAA, auxin-action inhibitor PCIB, and auxin-transport inhibitor NPA on seedling growth of TNG67 and IR64 during AG. Seeds were germinated and grown under fully embedded conditions for 10 days at 28*°*C in a 16-h light (5,000 lux)/8-h dark cycle. After sterilization, seeds were placed on 20 mL of 0.025% (w/v) Hyponex No.2 medium solidified with 4 mM CaCl_2_ and 0.25% phytagel (Sigma) and supplemented with IAA, PCIB, and NPA at indicated concentration in the test tube. The test tube was then filled with 70 mL of sterile water containing 4 mM CaCl_2_ and 2×10^-2^ g/L phytagel. Five milliliters of the solid medium were used to embed the seeds to avoid seeds from floating. Greening rate, rooting rate, shooting rate, seminal root length, and seedling height were measured daily and data were presented in Supplemental Figure S3. (A) Morphology of TNG67 seedlings at 5 DAI (left of the panel) and 8 DAI (right of the panel). (B) Morphology of IR64 seedlings at 5 DAI (left of the panel) and 8 DAI (right of the panel).

### MiR167 is an AG-responsive miRNA in rice

To determine whether auxin signaling and miRNAs are involved in response to AG tolerance in rice, seeds from TNG67 were germinated and grown in the dark for 6 days under aerobic or submerged conditions, and the small RNAs were purified from aerobic and hypoxic coleoptile and subjected to miRNA microarray analysis. As shown in Figure 5A, among 460 rice miRNAs presented on the array, the expression of most miRNAs was down-regulated in hypoxic coleoptile. Of those differentially expressed miRNAs, the expression levels of miR167 family were much higher in aerobic coleoptile than in hypoxic coleoptile (Figure 5, A and B). The miR167 family consists of 10 members in the rice genome and there is only one nucleotide difference in the 3’-end between miR167a-c and miR167d-j (Figure 5A). The expression of miR167a and miR167d was validated by stem-loop RT-qPCR and both miR167a and miR167d were expressed at significantly high levels in aerobic coleoptile but their expression was strongly suppressed underwater (Figure 5C). The small RNA gel blot analysis showed that the expression of miR167 was barely detectable in the hypoxic coleoptile but was significantly more present in the aerobic coleoptile of rice (Figure 5D).

**Figure 5.**
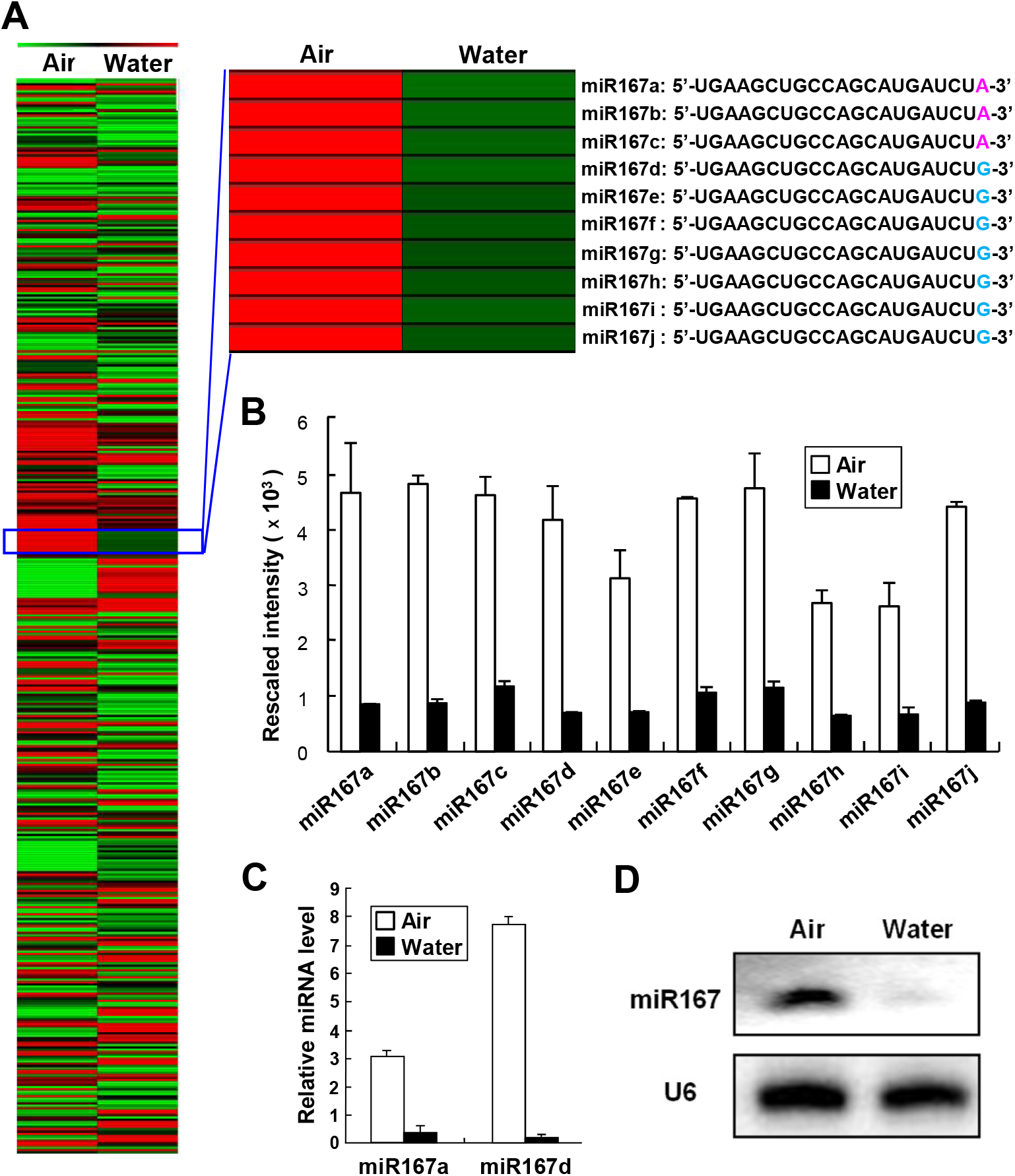
The Clustering of differentially expressed miRNAs in aerobic and hypoxic coleoptile. Rice seeds (*Oryza sativa* L. cv Tainung 67) were germinated and grown under aerobic or submerged conditions for 6 days at 28*°*C in the dark, and coleoptiles were harvested for small RNA extraction. The small RNAs were then subjected to microRNA array and clustering assay. Red and green indicate up-and down-regulated expression, respectively. (A) Four hundred sixty rice miRNAs are differentially expressed in aerobic (Air) and hypoxic (Water) coleoptiles. The expression of the miR167 family is up-regulated in aerobic coleoptiles. Sequence alignment of the miR167 family members is shown on the right of the panel. (B) Ten members of the miR167 family were expressed at a significantly higher level in aerobic coleoptile than in hypoxic coleoptile. (C) Expression of miR167a and miR167d in aerobic and hypoxic coleoptile by stem-loop RT-qPCR analysis. Rice coleoptile was grown for 6 days as described above. Small RNA was isolated from coleoptile and subjected to stem-loop RT reactions and qPCR analysis. (D) Small RNA gel blot analysis of miR167 expression in aerobic and hypoxic coleoptile. Total RNAs were purified from 6-day-old aerobic and hypoxic coleoptile and subjected to RNA gel blot analysis. Twenty micrograms of total RNAs were electrophoresed in 15% polyacrylamide gels and probed with γ-^32^P-labeled miR167 antisense. The expression of U6 snRNA was used as an internal control. *Error bars* represent the S.E. One-way ANOVA was used to analyze statistical difference; P < 0.01.

MiR167 was shown to be involved in the auxin regulation via the miR167-ARF-GH3 pathway in response to auxin signaling (Yang et al., 2006). To determine whether miR167 played a role in response to auxin signals and submergence during AG, the sequences of miR167a precursor and the artificial target-mimic of miR167, designated as MIM167, were fused downstream of the *Ubi* promoter (Figure 6A) and introduced into the rice genome. Several independent transgenic rice lines carrying the *Ubi::miR167a* were obtained, and three randomly selected lines were subjected to the small RNA gel bot assay. As shown in Figure 6B, the accumulation of miR167 was higher in the transgenic rice coleoptile than that in the wild-type in the hypoxic coleoptile. Only one transgenic rice line carrying the *Ubi:: MIM167* was obtained due to an extremely low transformation efficiency. The level of miR167 was reduced in the MIM167 transgenic rice coleoptile, particularly in the coleoptile grown in the air, compared to the coleoptile of wild-type rice (Figure 6C).

**Figure 6.**
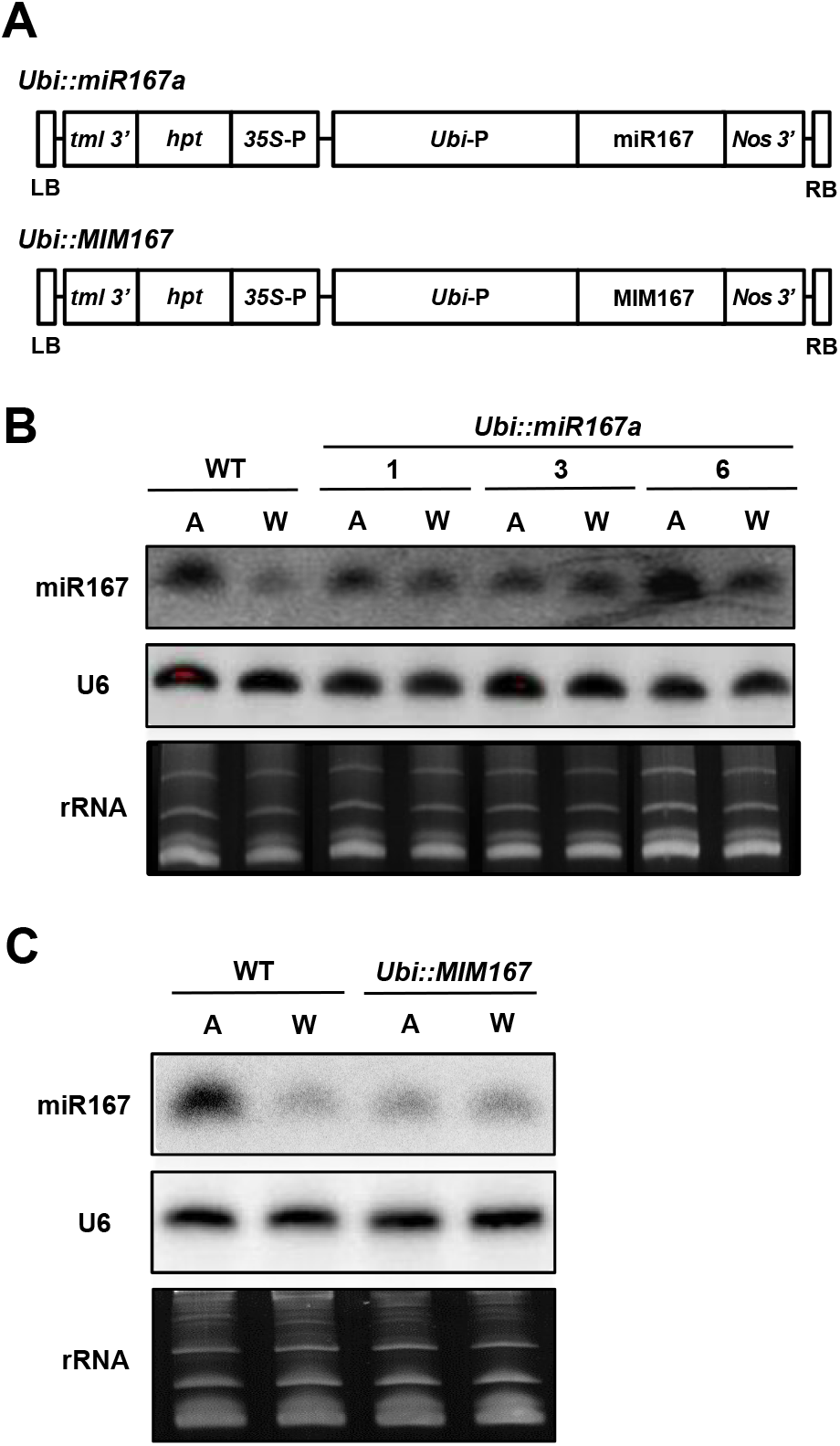
The expression level of miR167 in miR167a overexpression and target-mimic transgenic lines grown in aerobic and hypoxic condition. (A) Schematic representation of constructs of the binary vector used for rice transformation. Expression of miR167 in transgenic rice lines miR167a-1, miR167a-3, and miR167a-6 carrying the *Ubi::miR167a* gene (B), and miR167 target-mimic line MIM167 carrying the *Ubi::MIM167* gene (C). Seeds from the wild type and T2 seeds of homozygous transgenic rice were germinated and grown under aerobic or submerged conditions for 6 days. Total RNA was purified from aerobic (A) and hypoxic (W) coleoptile and subjected to RNA gel blot analysis as described in Figure 5D. The expression level of U6 snRNA was used as an internal control. Equal RNA sample loading was demonstrated by ethidium bromide staining of rRNA.

Four *OsARF* genes, *OsARF6*, *OsARF12*, *OsARF17*, and *OsARF25*, were reported to be the target genes of miR167 (Liu et al., 2012). It was shown that miR167-directed *OsARFs* mRNA cleavage is accompanied by down-regulation of the *OsGH3* genes (Yang et al., 2006; Ding et al., 2008). Overexpression of miR167a in the transgenic coleoptile grown underwater resulted in a significant decrease in mRNA levels of these four *OsARF* genes, *OsGH3-6*, and *OsGH3-8* (Supplemental Figure S4A). On the contrary, reduced miR167 level in aerobic coleoptile of the transgenic target-mimic line MIM167 displayed increased mRNA accumulation of these downstream-regulated genes (Supplemental Figure S4B).

### Reduced miR167 level results in enhanced tolerance to submergence during AG

To determine whether the expression level of miR167 could affect the accumulation of endogenous free IAA and the growth of rice, wild-type rice, miR167a overexpression transgenic rice lines, and target-mimic transgenic rice line MIM167 were grown and analyzed. It was found that, compared to WT, reduced miR167 level in the ten-day-old seedlings of transgenic rice line MIM16 in the air caused a substantial decrease in the endogenous accumulation of IAA. In contrast, no significant difference in miR167 level was observed among miR167a overexpression lines and WT (Supplemental Figure S5A). The target-mimic line MIM167 had a shorter shoot, but the seminal root length was longer than the WT and miR167a overexpression lines (Supplemental Figure S5, B to E). After transplantation to the paddy field, the MIM167 produced much more tiller numbers than the WT and miR167a (Supplemental Figure S6, A and C). The plant of miR167a was taller than WT, whereas the MIM167 plant exhibited the phenotype of semidwarf, with a 21% decrease in height than the WT (Supplemental Figure S6B). The panicle length of MIM167 was shorter than the WT and the grain number of the miR167a and MIM167 was significantly lower than the WT (Supplemental Figure S6, D, E and F). The thousand grains weight of the MIM167 was lower and the seed size was slightly smaller than the WT and miR167a (Supplemental Figure S6, G and H). The level of free IAA from freshly harvested grains was quantified and miR167a transgenic plants had the highest free IAA level while MIM167 transgenic plants had the lowest level of free IAA (Supplemental Figure S6I). In the paddy field, reduced miR167 level in adult plants resulted in noticeable phenotypic changes, particularly in inflorescence architecture and development (Supplemental Figure S6, F and J).

The target-mimic transgenic line MIM167 had reduced level of miR167, resulting in significantly decreased accumulation of endogenous free IAA and changes in the growth of transgenic rice plants. To determine whether MIM167 had enhanced tolerance to submergence during AG, seeds from WT, miR167a overexpression, and target-mimic MIM167 rice lines were germinated and grown under fully embedded conditions for 13 days in a light-dark cycle. The seeds were placed in a test tube filled with sterile water containing 2.0, 2.5, 3.0, and 4.0 x10^-2^ g/L phytagel. As shown in Figure 7, when the concentration of phytagel was 2.0-2.5 g/L, the MIM167 had better root and primary leaf growth than the WT seedlings, whereas the miR167a exhibited relatively poor seedling growth at 5, 7, and 9 DAI. When the concentration of phytagel was 3.0-4.0 g/L, hypoxia became more intense and the coleoptile elongated and overgrew in all three rice seedlings at 5, 7, and 9 DAI, and at 13 DAI, the root and primary leaf failed to grow in the WT and miR167a seedlings and only the MIM167 produced root and primary leaf. At 2.0 x10^-2^ g/L phytagel, the rate of greening, rooting, shooting, seminal root length, and seedling height of the MIM167 seedlings were higher than those of the WT and miR167a seedlings (Supplemental Figure S7). These results indicate that reduced miR167 level can enhance the tolerance of transgenic rice hypoxic seedlings to submergence during AG.

**Figure 7.**
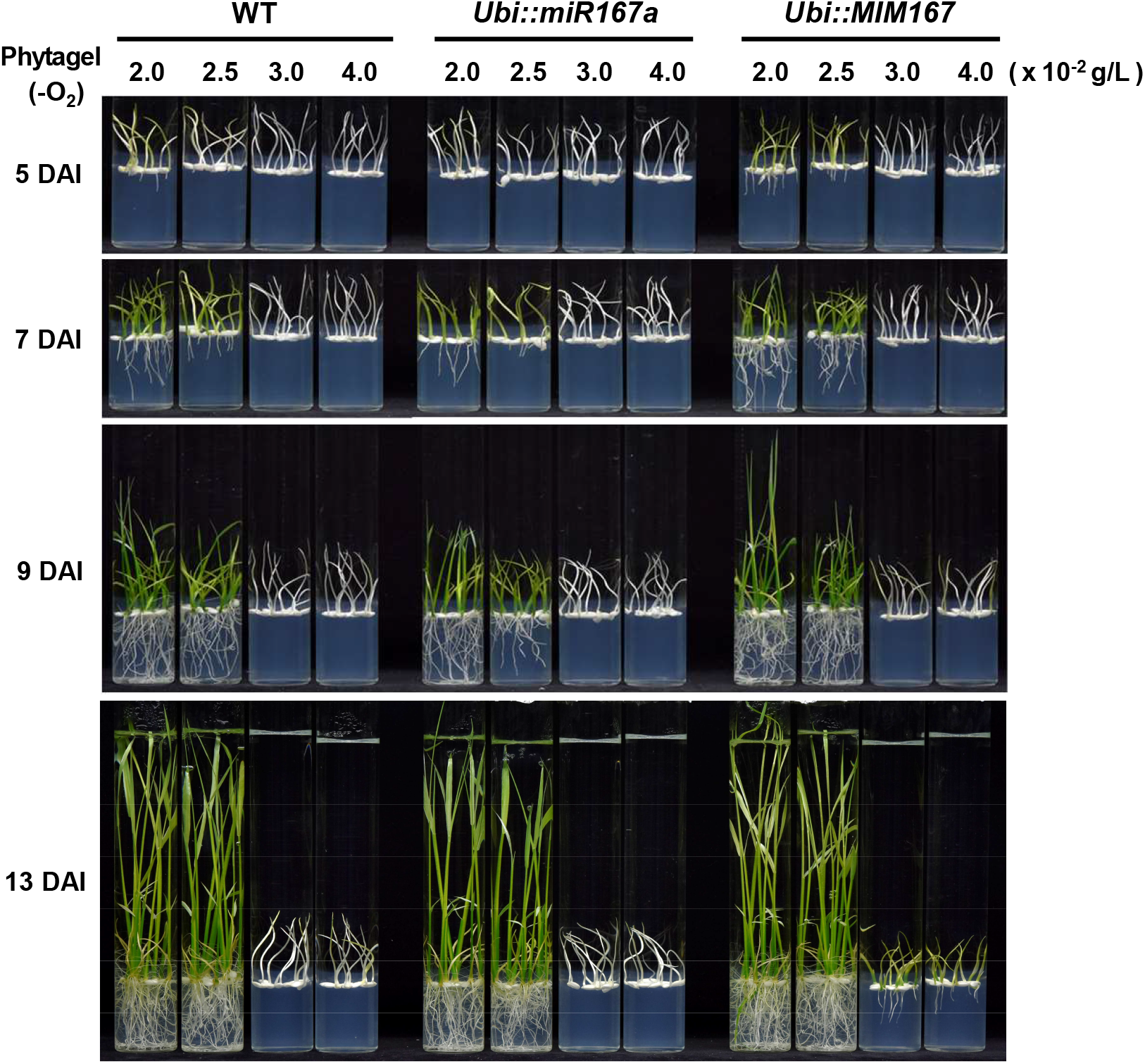
The miR167 target-mimic line MIIM167 exhibits improved tolerance to submergence compared to wild type and miR167a overexpression line during AG. Seeds from wild type (WT), transgenic rice lines miR167a and MIM167 were germinated and grown under fully embedded conditions for 13 days, as described in Supplemental Figure S1. The test tube was filled with 70 mL of sterile water containing 2.0, 2.5, 3.0, and 4.0 x10^-2^ g/L phytagel, respectively, and images of seedling morphology of WT, miR167a, and MIM167 at 5, 7, 9, and 13 DAI were shown. Greening rate, rooting rate, shooting rate, seminal root length, and seedling height at each time point were measured and shown in Supplemental Figure S7.

### Overexpression of *OsGH3-8* significantly promotes the seedling establishment and enhances the AG tolerance

We have demonstrated that AG-susceptible rice variety IR64 had high-level accumulation of IAA in hypoxic seedlings and these seedlings exhibited poor seedling establishment and reduced AG tolerance (Figure 3). The expression levels of genes related to auxin homeostasis, including miR167 and its downstream-regulated genes and auxin biosynthesis, signaling, and metabolism genes in TNG67 and IR64 seedlings were analyzed. Seeds were germinated and grown in a light-dark cycle under aerobic or submerged conditions for four days and seedlings were harvested and the embryo-coleoptile tissues were collected. Total RNA was purified from the collected embryo-coleoptile tissues and subjected to stem-loop RT-qPCR and quantitative real-time RT-PCR analysis. As shown in Figure 8, higher expression level of miR167a and miR167d was detected in seedlings grown under aerobic conditions for both TNG67 and IR64 while both miRNA167s were expressed at lower levels in IR64 than TNG67. As being the target genes of miR167, the expression levels of *ARF6* and *ARF17* inversely correlated with miR167a and miR167d levels but *ARF12* expression level was similar to that of both miR167s in both TNG67 and IR64; opposite results were obtained regarding the expression level of *ARF25* in seedlings grown under aerobic and submerged conditions between TNG67 and IR64. The expression of three auxin-responsive genes encoding IAA-amido synthetase, *GH3-2*, *GH3-6*, and *GH3-8*, was highly induced in seedlings grown underwater. Among them, the expression level of *GH3-6* and *GH3-8* correlated with that of *ARF6* and *ARF17* but inversely correlated with that of miR167s, confirming the involvement of miR167-ARF-GH3 pathway. The expression levels of *IAR3*, an IAA-amino acid hydrolases gene and one of the target genes of miR167 (Kinoshita et al., 2012), and three auxin biosynthetic genes, *YUCCA2*, *YUCCA3*, and *YUCCA*6, were also significantly higher in hypoxic seedlings of both TNG67 and IR64. In contrast, the expression pattern of another auxin biosynthetic gene, *TAA1*, was similar to that observed with *ARF25*. The expression profile of *IAR3* transcripts did not correlate with that of both miR167s in TNG67 and IR64, suggesting that expression of *IAR3* gene is not merely regulated by miR167s. These results suggested that high-level free IAA accumulation in hypoxic rice seedlings, especially in IR64, involved metabolic regulation of IAA biogenesis, which could be mediated through the miR167a-ARF-GH3 pathway. In addition, high-level expression of *GH3* genes might be required for the regulation of endogenous IAA levels and promotion of auxin homeostasis in rice in response to submergence.

**Figure 8.**
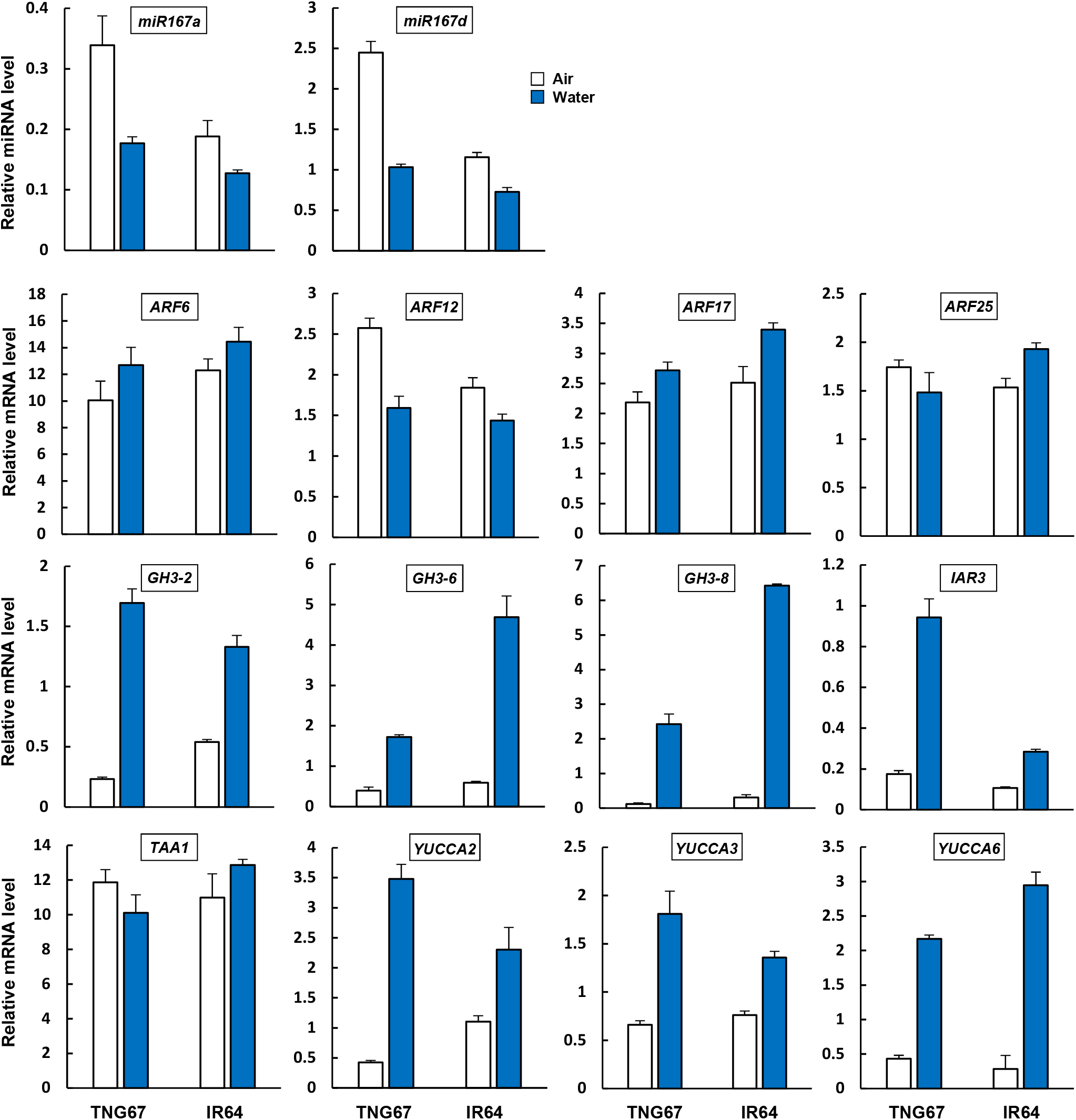
Expression analyses of miR167a, miR167d, auxin biosynthetic, responsive, and metabolic genes in the embryo-coleoptile tissues from TNG67 and IR64 rice varieties. Rice seeds of TNG67 and IR64 varieties were germinated and grown in aerobic (Air, *open bar*) or submerged (Water, *blue bar*) conditions as described in Fig. 3. Four-day-old aerobic and hypoxic seedlings were harvested and the embryo-coleoptile tissues were collected. Total RNA was purified from the collected embryo-coleoptile tissues and subjected to stem-loop RT-qPCR and quantitative real-time RT-PCR analysis. *Error bars* represent the S.E. of three replicates. Auxin response factors: *ARF6*, *ARF12*, *ARF17*, and *ARF25*; IAA-amido synthetases: *GH3-2*, *GH3-6*, and *GH3-8*; IAA-amino acid hydrolases: *IAR3*; IAA biosynthetic genes: *TAA1*, *YUCCA2*, *YUCCA3*, and *YUCCA6*. The expression level of eukaryotic elongation factor 1-alpha (*eEF1α*) was measured and served as the internal control for normalization. Accession numbers of indicated genes are provided in Supplemental Table S1.

Previous studies have shown that overexpression of *OsGH3-8* under the control of a constitutive promoter leads to a significant decrease in IAA level, which causes abnormal plant morphology and retarded growth and development in rice (Ding et al., 2008). To determine whether increased *OsGH3-8* level could directly promote AG tolerance in rice, a highly inducible promoter under hypoxia, *mαAmy8* promoter (Wu et al., 2014b), was used to express the *OsGH3-8* gene (Supplemental Figure S8A). The *mαAmy8::OsGH3-8* chimeric gene was introduced into the rice genome via *Agrobacterium*-mediated transformation. Several independent transgenic rice lines were obtained, and seeds were germinated and grown in the air or under fully embedded conditions as described previously. We found that overexpression of *OsGH3-8* in transgenic seedlings grown in the air results in greatly enhanced growth of seminal root at 4 and 5 DAI compared to the WT seedlings (Supplemental Figure S8, B to D), suggesting that such emhancement may be resulted from decreased endogenous IAA level in transgenic rice seedlings. In transgenic hypoxic seedlings, increased *OsGH3-8* levels significantly promoted seedling establishment and enhanced the AG tolerance in transgenic rice lines at 9 and 13 DAI, particularly as the phytagel concentration was 3.0-4.0 x 10^-2^ g/L (Figure 9 and Supplemental Figure S9). In addition. Seedlings of transgenic rice lines carrying the *mαAmy8::OsGH3-8* gene also displayed greater submergence tolerance than TNG67 and IR64 seedlings at 15 DAI even the concentration of phytagel was at 2.5 g/L (Supplemental Figure S10).

**Figure 9.**
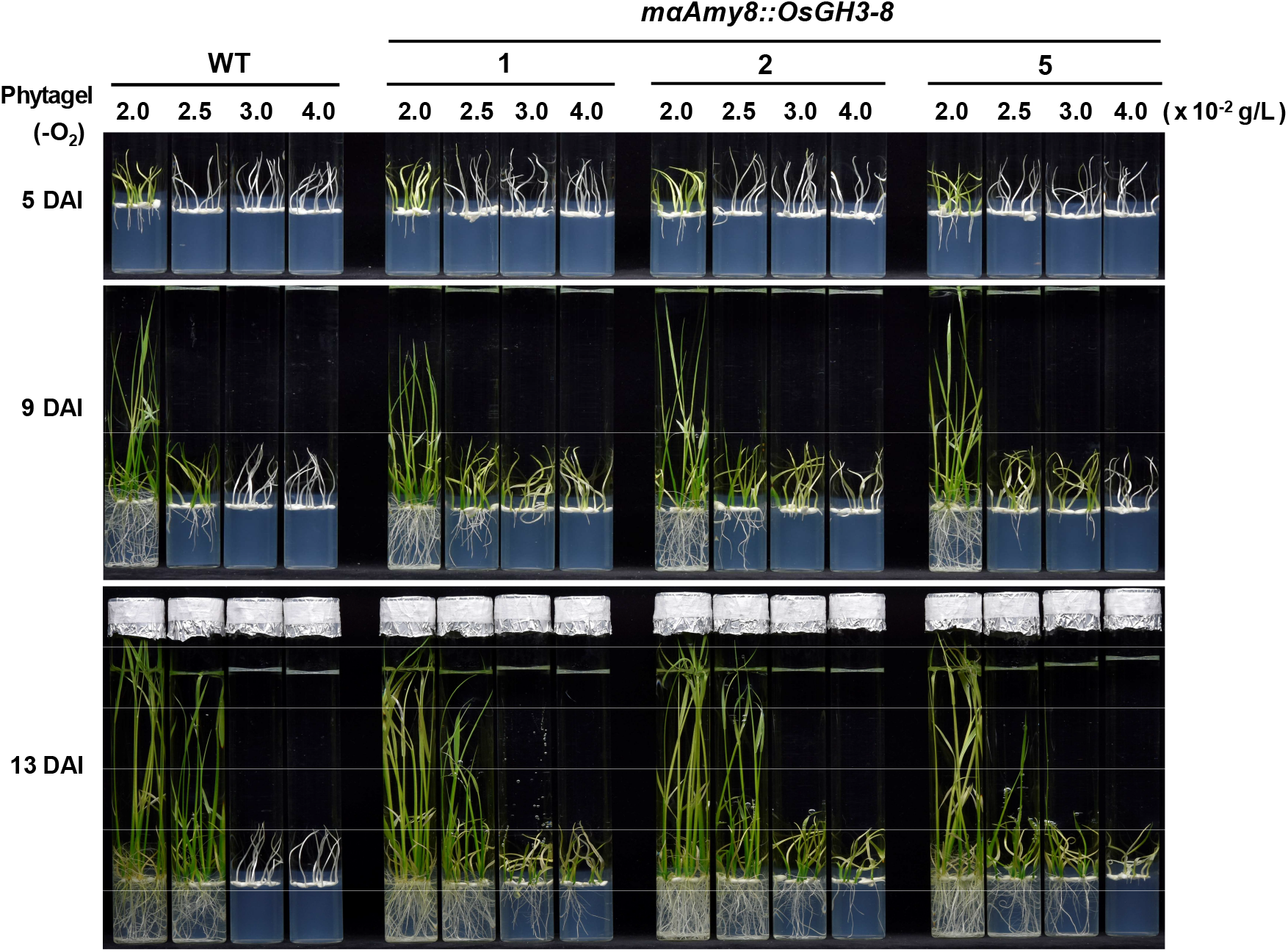
Overexpression of *OsGH3-8* under the control of the modified *αAmy8* promoter in hypoxic seedlings confers enhanced tolerance to submergence during AG. T2 seeds of homozygous transgenic rice lines 1, 2, and 5 carrying the *mαAmy8::OsGH3-8* gene were obtained. Seeds from WT and transgenic rice were germinated and grown under fully embedded conditions, as described in Figure 7 and images of seedlings at 5, 9, and 13 DAI were shown. Greening rate, rooting rate, shooting rate, seminal root length, and seedling height at each time point were measured and shown in Supplemental Figure S9.

## Discussion

### Auxin-mediated growth inhibition in hypoxic seedlings depends on high endogenous levels of IAA

The phytohormone auxin has long been known to be important in stimulating coleoptile elongation and rapid seedling growth in the air (Gallei et al., 2020), but little is known regarding its role in influencing the elongation of rice coleoptile and the production of root and primary leaf underwater. To understand auxin’s role in rice under anoxic and hypoxic conditions, the phototropic responses of rice seedlings under submergence was analyzed. We found that hypoxic coleoptile was positively phototropic, whereas the root showed negative phototropism regardless of whether seedlings were grown in the air or underwater (Supplemental Figure S11). Under submergence, the accumulation of free IAA in coleoptile stimulates its elongation. Conversely, higher IAA levels on the shaded side lead to root growth inhibition, which is similar to root growth inhibition in the air. It suggests that rice seedlings are sensitive to endogenous auxin under submergence. A recent report has also demonstrated that auxin is required for rice seed germination underwater; their results indicated that auxin availability and transport play a critical role in determining the final coleoptile length in *japonica* rice (Nghi et al., 2021). However, the role of auxin in the regulation of rice adaptation to AG and seedling establishment is still unclear.

When rice seeds were germinated in submerged fields, several environmental factors could affect seed germination and early seedling growth in rice. Based on the water status, the dissolved oxygen concentration in water varies in different ecological conditions from < 0.003 mol O_2_ m^-3^ (anoxia) or 0.03 mol O_2_ m^-3^ (hypoxia) to 0.25 mol O_2_ m^-3^ (aerated water)(Ismail et al., 2009). Therefore, hypoxic rice seedlings experience different intensities of low oxygen tension during AG and seedling growth. In submerged fields, rapid coleoptile emergence results in the appearance of a coleoptile tip above the soil during AG; meanwhile, coleoptile elongation and seedling establishment were affected by light (Biswas and Yamauchi, 1997). Light is known to control various physiological processes in the plant, such as leaf expansion, shade-avoidance response, seed germination, the greening of seedlings, and chloroplast development (Salazar-Iribe and De-la-Peña, 2020; Cackett et al., 2021; Wang et al., 2022). It has been demonstrated that phytochrome B (phyB) is involved in these processes by regulating auxin levels in response to light conditions (Zhang et al., 2014). Under low R: FR conditions or in darkness, phyB exists mainly in the inactive Pr form, allowing the accumulation of phytochrome-interacting factors (PIFs) to promote the expression of auxin biosynthetic genes, transporter, and auxin response genes (Li et al., 2012). Conversely, phyB exists mainly in the active Pfr form in high R: FR conditions, which modulates the activation of *SUPERROOT* (*SUR2*) expression to decrease auxin level by repressing auxin biosynthesis and promoting auxin homeostasis (Delarue et al., 1998; Hoecker et al., 2004).

In the present study, we found that under dark submergence, IAA levels in hypoxic rice seedlings grown under stagnant water were high enough to repress root and primary leaf emergence but stimulate coleoptile elongation (Figure 1). Conversely, oxygen and light significantly reduce the endogenous levels of free IAA and promote root and primary leaf growth under submergence (Figures 1 and 2). Comparison of AG-tolerant and AG–intolerant rice varieties reveals that intolerant variety IR64 has a poor seedling establishment and relatively higher accumulation of endogenous IAA in hypoxic seedlings during AG (Figure 3). According to the results of exogenous PCIB treatment, we found that reducing the auxin action significantly enhanced seedling establishment and AG tolerance in TNG67 and IR64 hypoxic seedlings (Figure 4). Overall, these results suggest that seedling growth inhibition in response to excessive IAA levels would cause intolerance to submergence, and a certain threshold level of IAA may be required for AG tolerance in rice. In previous studies, *DR5:GUS* was used to identify the site of auxin response in germinated rice seeds and a higher accumulation of auxin at the vasculature tissues of the scutella, coleoptile, primary leaf, and in the root stele, root cap, and endosperm was detected (Guo et al., 2016). Whether or not the auxin transport or distribution in hypoxic rice seedlings would be influenced by excessive accumulation of endogenous free IAA and affect the AG tolerance remains to be explored.

### The miR167-ARF-GH3 pathway is essential in response to auxin signaling and AG tolerance

MicroRNA expression is highly sensitive to environmental stresses that function as essential regulators in adapting biotic and abiotic stresses with plant development (Rubio-Somoza and Weigel, 2011). Previous studies have demonstrated that miR167 can inhibit the expression of *ARF6, ARF7,* and *IAR3* genes in *Arabidopsis* and plays a critical role in regulating the development of the flower, adventitious root, and lateral root in response to auxin signaling (Wu et al., 2006; Gutierrez et al., 2009; Kinoshita et al., 2012; Jodder, 2020), and may be involved in the control of tiller angle in rice (Li et al., 2020). Previous studies have also demonstrated that the miR167-ARF-GH3 pathway is involved in response to exogenous application of auxin in cultured rice cells (Yang et al., 2006), hinting at the role of miR167 in the regulation of endogenous IAA levels and the maintenance of auxin homeostasis by activating *GH3* genes expression. In this study, high-level accumulation of miR167a and miR167d was detected in coleoptile and embryo of aerobic rice seedlings, whereas their expression is significantly repressed under submergence (Figures 5 and 8). In the target-mimic rice line MIM167, reduced miR167a levels lead to a substantial decrease in the accumulation of endogenous free IAA (Supplemental Figures S5A and S6I) and consequently promote the seedling establishment and enhance the AG tolerance (Figure 7). A similar result was observed in the transgenic rice line carrying the *mαAmy8::OsGH3-8* chimeric gene (Figure 9), *OsGH3-8* is highly expressed in hypoxic rice seedlings, and it exhibits high tolerance to submergence (Figure 9 and Supplemental Figure S10). These results indicate that the miR167-ARF-GH3 pathway is involved in the metabolic regulation of IAA, and possibly required for AG tolerance in rice.

As shown in Figures 3B and 8, the accumulation of miR167a and miR167d in TNG67 and IR64 seedlings grown in the air or under submergence is inversely correlated with endogenous IAA levels, suggesting that the expression of miR167 is not merely due to submergence repression but might be regulated by feedback inhibition of endogenous IAA. Thus, it was proposed that endogenous free IAA levels modulate seedling establishment of rice during AG through the miR167-ARF-GH3 pathway. As shown in Figure 10, oxygen deprivation under dark submergence promotes the expression of auxin biosynthetic genes (*YUCCA2*, *YUCCA3*, and *YUCCA6*), leading to an excessive accumulation of endogenous free IAA in hypoxic rice seedlings, which stimulates coleoptile elongation but represses root and primary leaf formation. Conversely, oxygen and light reduce the accumulation of free IAA, promote seedling establishment, and enhance AG tolerance. The expression of miR167 is repressed by submergence and possibly feedback suppression of endogenous IAA to modulate free IAA accumulation through the miR167-ARF-GH3 pathway. Reduced miR167 level prevents *ARF* mRNA cleavage and subsequently activates the *GH3* gene expression, increasing auxin metabolism and promoting auxin homeostasis. In addition, due to reduced miR167 expression by submergence, *IAR3* expression increased, leading to excessive auxin levels in hypoxic seedlings as a result of increasing hydrolysis of IAA-Ala back to free IAA via IAR3 hydrolase.

**Figure 10.**
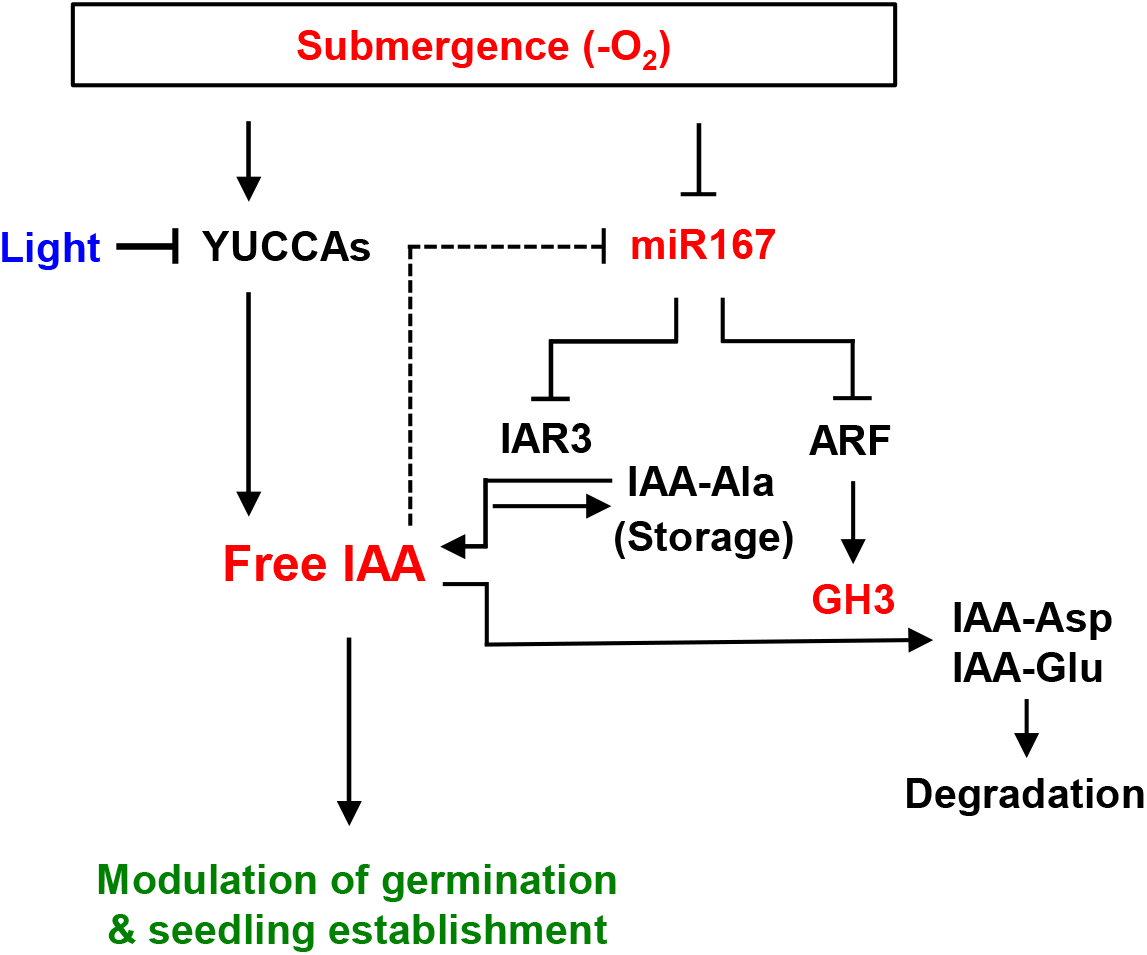
The molecular mechanisms involved in the interplay between oxygen, light, and IAA signaling modulate seed germination and seedling establishment under submergence in rice.

### The greening of hypoxic coleoptile is vital in response to AG tolerance

It was previously shown that carbohydrate availability played a pivotal role in survival to submergence in rice (Loreti et al., 2018); however, under dark submergence, the addition of sugar results in promoting coleoptile elongation and the inability to enhance seedling establishment during AG (Li et al., 2014). Therefore, another critical regulatory factor might affect rice seedling establishment and AG tolerance. In the present study, it was demonstrated that endogenous IAA levels could modulate rice’s seedling establishment and AG tolerance under submergence. It was also found that oxygen and light reduced the excessive accumulation of endogenous free IAA, promoted seedling establishment, and enhanced AG tolerance (Figures 1 and 2), and the greening of the coleoptile plays a pivotal role in the subsequent production of the root and primary leaf under submergence (Supplemental Figure S1). As shown in Figure 4 and Supplemental Figure S3, the addition of IAA negatively reduces the greening rate of hypoxic seedlings, and the timing of the greening of coleoptile was delayed in TNG67 and IR64. On the contrary, earlier greening timing was observed when IAA function was inhibited, suggesting that the greening of the coleoptile in response to auxin signaling might play a vital role in influencing seed germination and seedling establishment under submergence in rice. It has been reported that chloroplast development and photomorphogenesis in green plant tissues depend on the interplay between light and phytohormones signaling pathways, including auxin signaling (Salazar-Iribe and De-la-Peña, 2020; Cackett et al., 2021; Wang et al., 2022) but the interplay underlying these signaling pathways in modulating AG tolerance is still unclear. Moreover, whether sugar could affect the accumulation of endogenous free IAA to influence AG and seedling establishment in rice remains elusive.

## Materials and methods

### Plant material

The rice cultivar used for all experiments in this study was *Oryza sativa* L. cv. Tainung 67 (TNG67). Seeds of *japonica* rice varieties Nipponbare (Nipp) and Taichung 65 (TC65) were provided by Taiwan Agricultural Research Institute (TARI), and seeds of four rice varieties, Khao Hlan On (KHO), IR42, IR64, and FR13A, were obtained from the International Rice Germplasm Center (IRGC) of the International Rice Research Institute (IRRI). Some rice varieties were used to measure endogenous IAA levels, real-time quantitative RT-PCR analysis, and phototropism assays. Seeds from TNG67 were sterilized with 3 % sodium hypochlorite for 30 min, washed extensively with sterile distilled water, and placed on N6D agar medium (Toki, 1997) for callus induction. After one month of culture, calli derived from scutellum were sub-cultured in a fresh N6D medium for transformation.

### Seed germination and seedling growth under aerobic and submerged conditions

Rice seeds were sterilized with 3% NaOCl and rinsed with sterile distilled water. For germination in air, seeds were placed on 20 mL of 0.025% (w/v) Hyponex No.2 medium solidified with 4 mM CaCl_2_ and 0.25% phytagel (Sigma) in the test tube. For germination under submerged conditions, seeds were placed in a test tube as in germination in the air experiment, and the test tube was filled with 70 mL of sterile water lacking phytagel. The water was either continuously bubbled with air (Aerated) or not circulated or refreshed (Stagnant) during the treatment. For germination under fully embedded conditions, the test tube was filled with 70 mL of sterile water and supplemented with 4 mM CaCl_2_ and phytagel at the indicated concentration in water. Five milliliters (mL) of the solid medium were used to embed the seeds during anaerobic germination to avoid seeds from floating. Subsequently, seeds were allowed to germinate and grow at 28°C in the dark or in a 16-h light (5,000 lux)/8-h dark cycle. Aerobic and hypoxic tissues were collected and assayed for subsequent experiments. Oxygen concentration in the water was determined by a dissolved oxygen meter (Model EcoScan D06, Eutech Instruments Pte. Ltd.). The O_2_ concentration was approximately 0.25 mol m^-3^ in aerated water and 0.1 mol m^-3^ in stagnant water.

### Quantification of IAA, IAA-amino acid conjugates, and oxIAA

The sample preparation for quantifying free IAA, IAA-amino acid conjugates, and oxIAA was performed according to the method described by Novák *et al.* (2012). Endogenous free IAA, conjugates, and metabolites were measured by ultra-performance liquid chromatography-electrospray ionization-tandem mass spectrometry (UPLC-ESI-MS/MS, Waters Xevo TQ-S). For quantification of IAA and its conjugates and metabolites, 600 mg of fresh rice tissues were extracted with extraction buffer (methanol:water: acetic acid, 80:19:1, v/v/v), and the stable isotope-labeled [^13^C_6_]-IAA (Cambridge Isotope Laboratories) was used as internal standard. Samples were purified by solid-phase extraction using Oasis^TM^ HLB extraction cartridge columns (Waters) as described previously (Novák et al., 2012). The purified samples were eluted with 2 mL of 80% methanol, the liquid was evaporated using a Speed Vac, and the residue was dissolved in 60 μL of 10% (v/v) methanol. The samples were then filtered through a 0.22 μm pore-sized PVDF membrane syringe filter (Jet Biofil) and subjected to UPLC-ESI-MS/MS analysis.

### Primers

All primers are listed in Supplemental Table S1 online.

### Plasmid construction and rice transformation

For the construction of plasmid containing the *Ubi::miR167a* chimeric gene, the precursor sequence of miR167a was amplified with the miR167a-F and miR167a-R primers using genomic DNA of rice (*Oryza sativa* L. cv. Tainung 67) as the DNA template. The PCR fragment containing 141-bp precursor sequence of miR167a was digested with *Bam*HI and then sub-cloned into pAHC18 (Bruce et al., 1989), containing *Ubi* promoter and *Luc* cDNA, to generate pUbi-miR167a.

For the construction of plasmid carrying the *Ubi::MIM167* chimeric gene, an artificial target-mimic sequence of miR167 (MIM167) was generated according to the method described by Franco-Zorrilla et al. (2007). First, a 76-bp DNA fragment containing the MIM167 was PCR-amplified with primers MIM167-F and MIM167-R by primer-dimer production. The MIM167 fragment was then sub-cloned into pAHC18 at *Bam*HI site, yielding pUbi-MIM167.

For the construction of plasmid containing the *mαAmy8::OsGH3-8* chimeric gene, plasmid pαAmy8(+G+2TA)-Ubi(in)-Luc containing the modified *αAmy8* promoter (Wu et al., 2014b) was used for high-level expression of *OsGH3-8* gene during AG. Plasmid pUbi-cmyc (Wu et al., 2014b) was digested with *Nco*I and *Bam*HI, and the restriction fragment containing a duplicated *c-Myc* epitope tag was then sub-cloned into the same site in pαAmy8(+G+2TA)-Ubi(in)-Luc to generate pmαAmy8-cmyc-Nos. The *OsGH3-8* gene was amplified from *OsGH3-8* cDNA (Accession No. AK101193) by PCR with the GH3-8-FW and GH3-8-RV primers. The 1829-bp DNA fragment containing the *OsGH3-8* coding region was flanked by the BamHI site and then sub-cloned into pmαAmy8-cmyc-Nos to generate pmαAmy8-cmyc-OsGH3-8-Nos.

To prepare the constructs for the *Agrobacteria*-mediated transformation, plasmids pUbi-miR167a and pmαAmy8-cmyc-OsGH3-8-Nos were linearized with *Hind*III and inserted into the corresponding site on binary vector pSMY1H (Ho et al., 2000), generating pAUbi-miR167a and pAm8-cmyc-OsGH3-8-Nos. Plasmid pUbi-MIM167 was digested with *Sph*I, blunt-ended, and inserted into the blunt-ended pSMY1H to yield pAUbi-MIM167. Plasmids pAUbi-miR167a, pAUbi-MIM167, and pAm8-cmyc-OsGH3-8-Nos were then introduced into *A. tumefaciens* strain EHA101 (Hood et al., 1986) with a MicroPulser electroporator (Bio-Rad) following the manufacturer’s instruction. Rice transformation was performed as described previously (Chen et al., 2002).

### Total and small RNA extraction

Total RNAs were purified from aerobic and hypoxic tissues of rice using TRIZol reagent (Invitrogen). The small RNAs were isolated using the PureLink™ miRNA Isolation Kit (K1570-01, Invitrogen) according to the manufacturer’s instructions.

### Rice miRNA microarray analysis

Rice miRNA microarray had been built in-house as described in a previous report (Liu et al., 2017). Rice seeds (*Oryza sativa* L. cv Tainung 67) were germinated and grown under aerobic or submerged conditions for 6 days at 28°C in the dark and the aerobic and hypoxic coleoptile were harvested for small RNA purification. Briefly, the purified miRNAs were labeled with Cy5 fluorophore (Cy5 Labeling Kit; Mirus Bio LLC) and then subjected to hybridization with the customized microarray for 4 hours in a 42°C oven. After hybridization and washing procedures, the microarray was scanned using a GenePix 4000B scanner, and the image was digitized by GenePix 4.0 software (Molecular Devices). Subsequently, the raw data of microarray were normalized using the quantile method, followed by a floor value assignment of 800 according to the recommendation of the chip manufacturer. The miRNAs with false discovery rate (FDR) < 0.05 and 2-fold change between the compared groups were identified as significantly differential genes.

### Small RNA gel blot analysis

Seeds from TNG67 and transgenic rice were germinated and grown under aerobic or submerged conditions for 6 days. Total RNA was purified from aerobic and hypoxic coleoptile and subjected to small RNA gel blot analysis. Twenty micrograms of total RNAs were separated on 15% acrylamide/8 M urea gel and transferred onto an Amersham™ Hybond™-N^+^ nylon membrane (GE Healthcare). The specific isotope probe was generated by end-labeling DNA oligonucleotides complementary to the miR167a sequence with [γ-^32^P] ATP by T4 polynucleotide kinase (New England Biolabs). The level of U6 snRNAs was used as the internal control. The hybridization was performed at 42°C for 16 hours in UltraHyb-oligo buffer (Ambion). The hybridized membrane was washed twice with washing buffer (2X SSC and 0.2% SDS) at 42°C for 3 min and then exposed onto an X-ray film.

### Semi-quantitative RT-PCR analysis

Semi-quantitative RT-PCR analysis was performed as described previously (Lee et al., 2009; Wu et al., 2014a). Seeds from TNG67 and transgenic rice were germinated and grown under aerobic or submerged conditions for 6 days. Total RNAs were purified from aerobic and hypoxic coleoptile and subjected to RT-PCR analysis. All amplifications were carried out with *Taq* DNA polymerase (Promega) using gene-specific primers (see Supplemental Information Table S1 online). RT-PCR amplification of the 192-bp amplicon of *TubA1* cDNA was performed as described previously (Wu et al., 2014a), and the level of *TubA1* mRNA was used as an internal control.

### Stem-loop quantitative RT-PCR and quantitative real-time RT-PCR analysis

To validate the differentially expressed miRNAs, the stem-loop quantitative reverse transcription PCR (stem-loop RT-qPCR) of candidates was performed as described previously (Varkonyi-Gasic et al., 2007). Total RNAs were reverse-transcribed into cDNAs using the miRNA-specific stem-loop RT primers. The expression level of miRNA was detected with SYBR Green reagent (FastStart SYBR Green Master, Roche) on a thermocycler. The expression level of eukaryotic elongation factor 1-alpha (*eEF1α*) served as the internal control (Jain et al., 2006b). The stem-loop RT-qPCR experiments were performed in triplicate. The detailed methodology and calculation of relative quantification were also described in the previous study (Liu et al., 2017).

To determine the expression level of genes related to IAA biosynthesis and metabolism, quantitative real-time RT-PCR was performed as described previously (Chen et al., 2006). Rice seeds of TNG67 and IR64 varieties were germinated and grown in aerobic or submerged conditions. Four-day-old aerobic and hypoxic seedlings were harvested, and the embryo-coleoptile tissues were collected. Total RNAs were purified from the collected embryo-coleoptile tissues and subjected to quantitative real-time RT-PCR. Real-time PCR was performed with a Rotor-Gene Q 2plex HRM thermocycler (Qiagen) using an SYBR Green reagent (Roche) and the amplification of *eEF1α* mRNA was used as an internal control for normalization. The primers for real-time PCR are listed in Supplemental Table S1 online.

## Supplemental data

The following materials are available in the online version of this article.

**Supplemental Figure S1.**
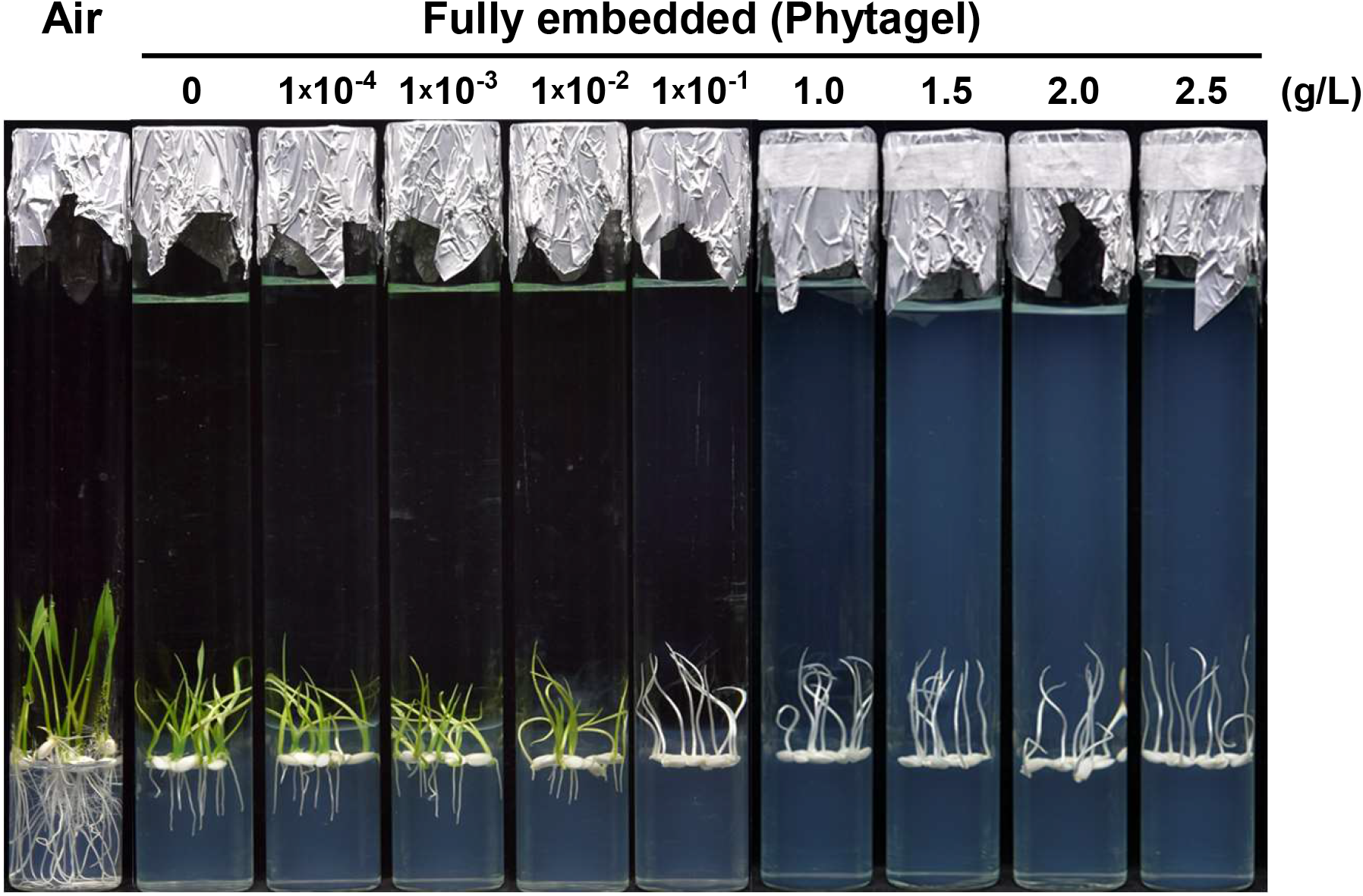
The oxygen level negatively affects seedling establishment under submergence. Rice seeds (*Oryza sativa* L. cv Tainung 67) were germinated and grown under aerobic conditions (Air) or fully embedded conditions for 5 days at 28*°*C in a 16-h light (5,000 lux)/8-h dark cycle. For germination in air, seeds were placed on 20 mL of 0.025% (w/v) Hyponex No.2 medium solidified with 4 mM CaCl_2_ and 0.25% phytagel (Sigma) in the test tube. For germination under fully embedded conditions, seeds were placed in a test tube as in germination in the air experiment and the test tube was filled with 70 mL of sterile water lacking or containing 4 mM CaCl_2_ and 1×10^-4^ to 2.5 g/L phytagel. Five milliliters of the solid medium were added to embed the seeds to avoid seeds from floating.

**Supplemental Figure S2.**
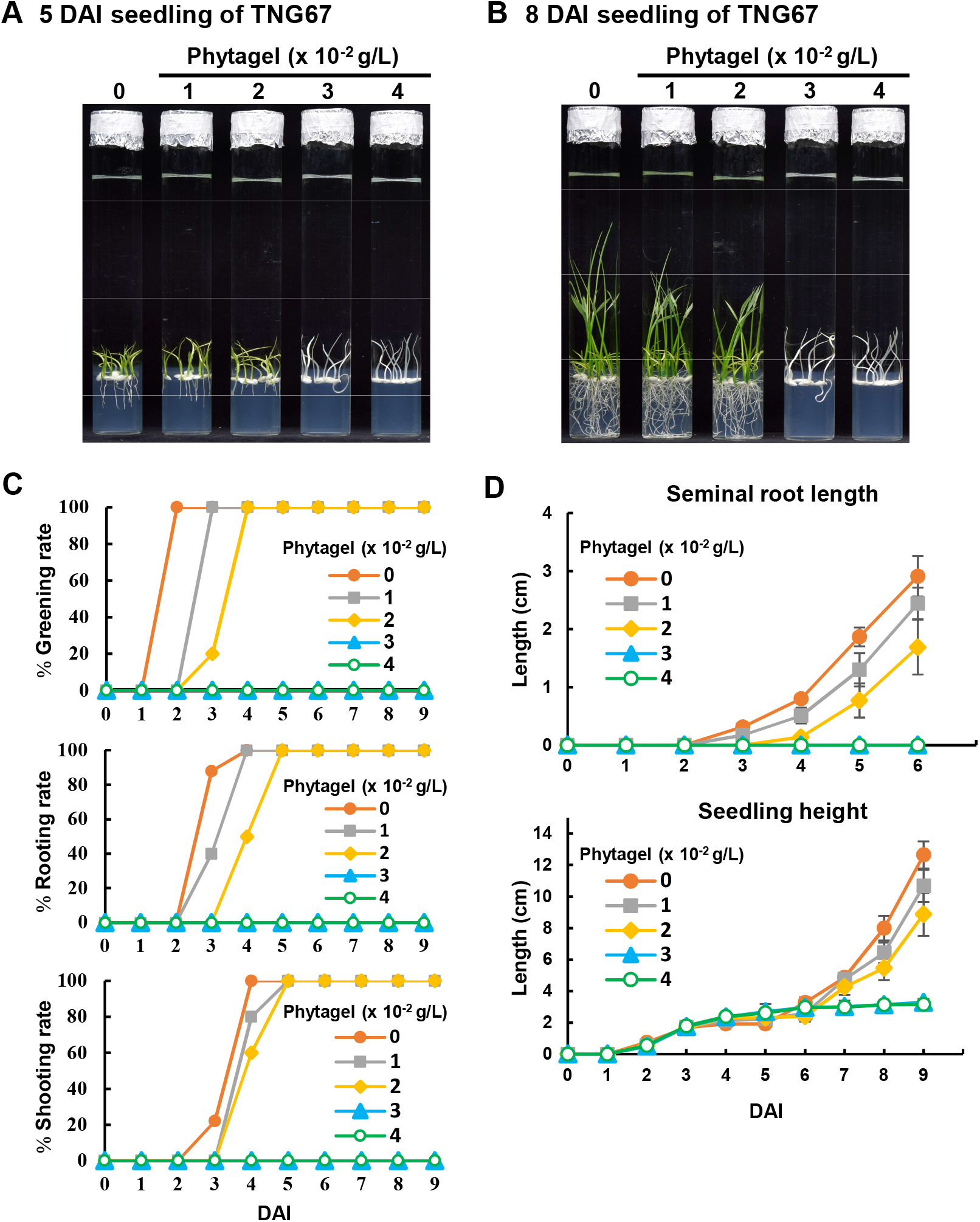
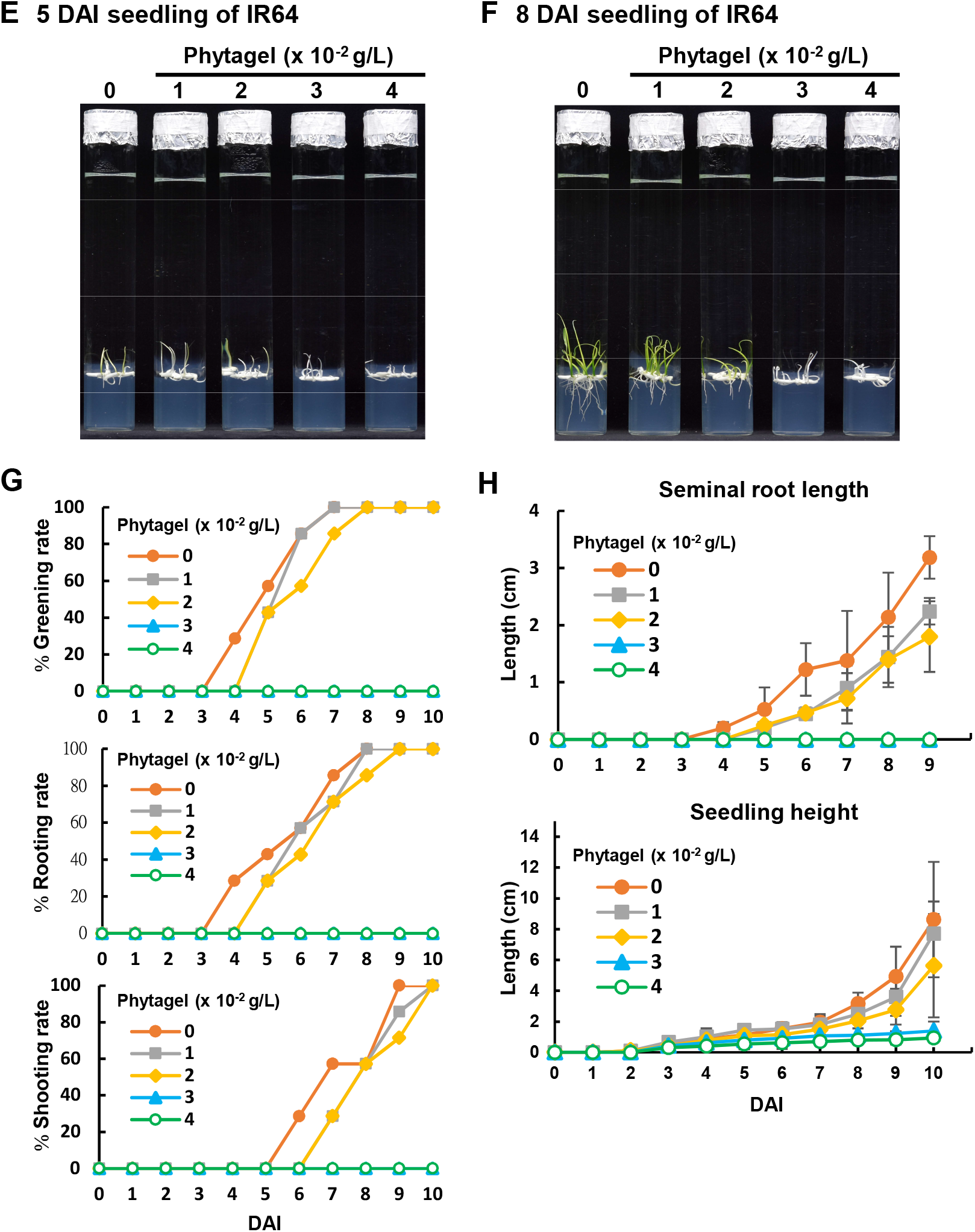
The rice cultivar TNG67 exhibits enhanced submergence tolerance than IR64 during AG. Seeds of *O. sativa* varieties TNG67 and IR64 were germinated and grown under fully embedded conditions as described in Figure S1. The test tube was filled with 70 mL of sterile water lacking or containing 4 mM CaCl_2_ and 1 to 4 x10^-2^ g/L phytagel. Greening rate, rooting rate, shooting rate, seminal root length, and seedling height were measured daily. (A) Morphology of TNG67 seedlings at 5 days after imbibition (DAI). (B) Morphology of TNG67 seedlings at 8 DAI. (C, D) Greening rate, rooting rate, shooting rate, seminal root length, and seedling height of TNG67 seedlings grown under fully embedded conditions. (E) Seedling morphology of IR64 at 5 DAI. (F) Morphology of IR64 seedling at 8 DAI. (G, H) Greening rate, rooting rate, shooting rate, seminal root length, and seedling height of IR64 seedlings grown under fully embedded conditions. *Error bars* indicate the S.E. of seedlings (n=10). Significance level was analyzed using One-way ANOVA; *P* < 0.01.

**Supplemental Figure S3.**
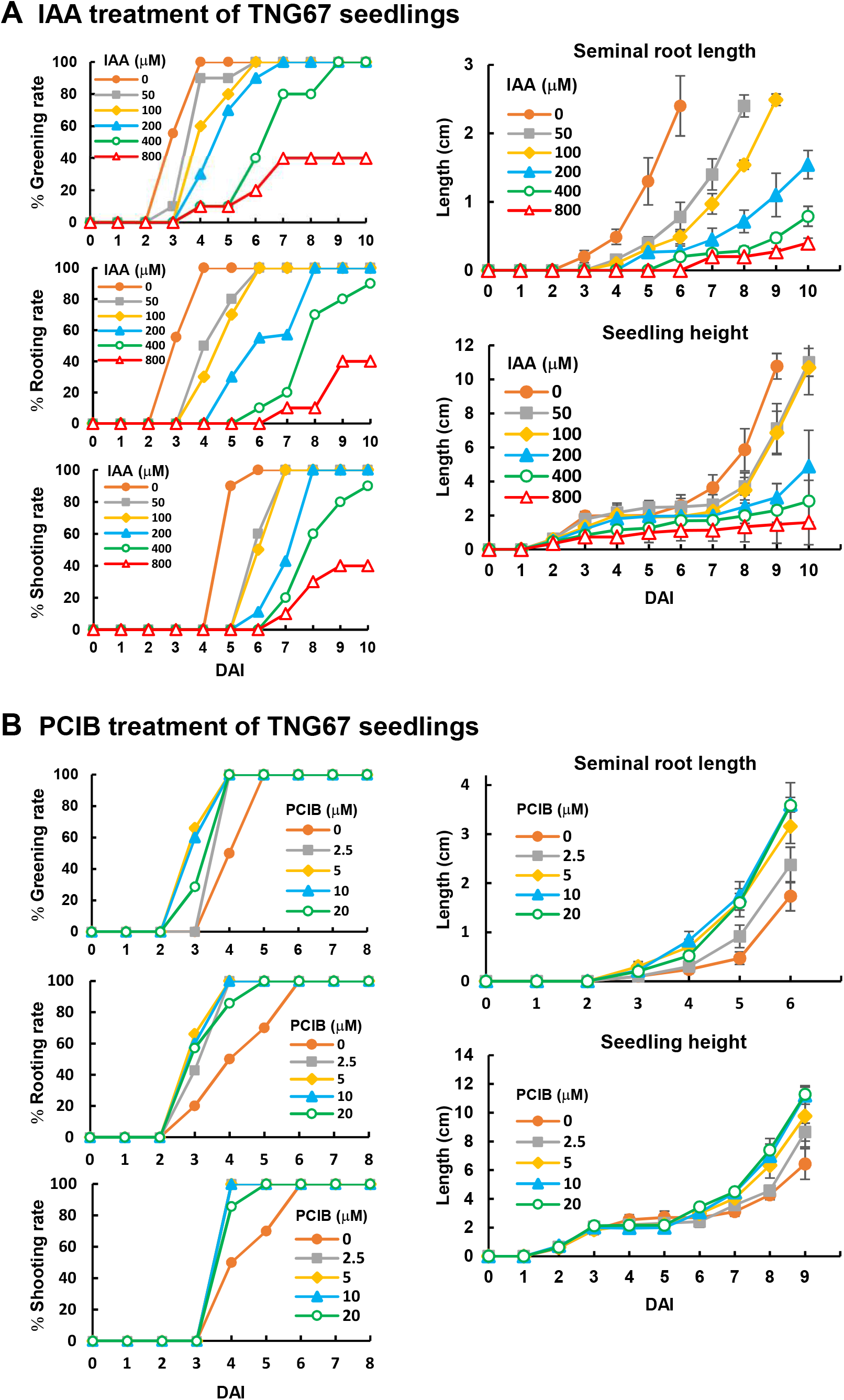
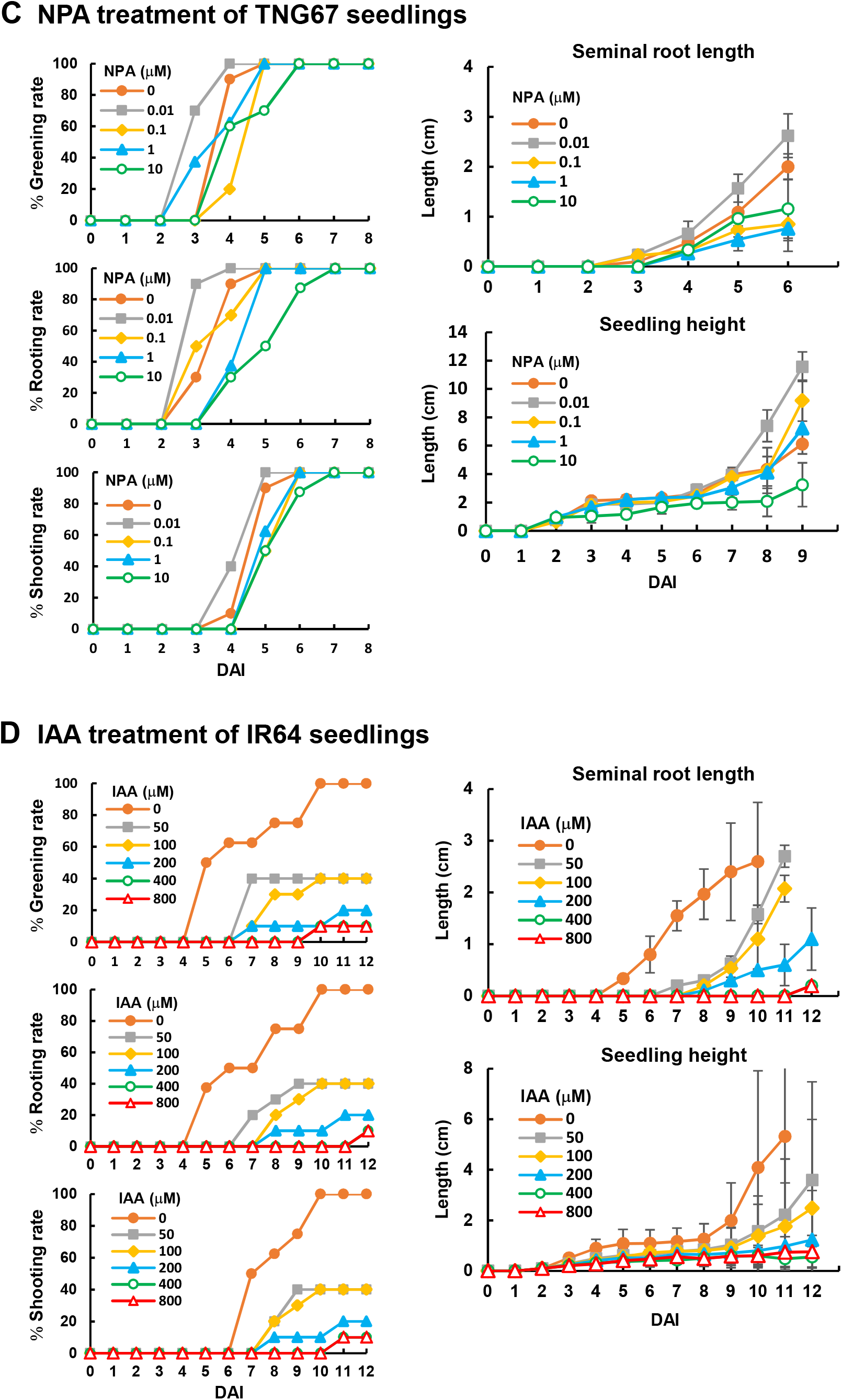
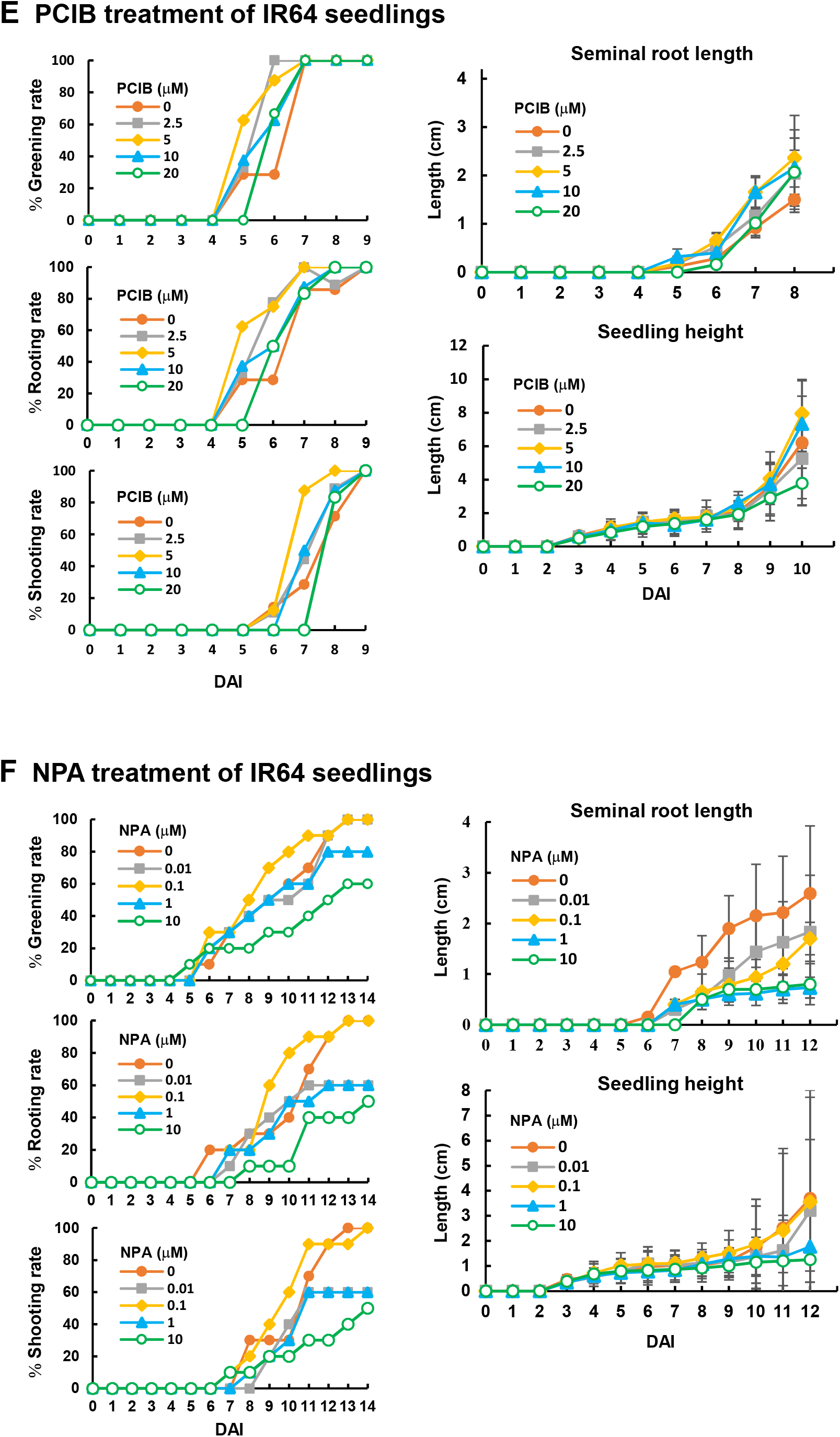
Effect of different concentrations of IAA, PCIB, and NPA on seedling growth of TNG67 and IR64 during AG. Seeds were germinated and grown under fully embedded conditions, as described in Figure 4. Greening rate, rooting rate, shooting rate, seminal root length, and seedling height were measured daily. Growth establishment of TNG67 seedlings treated with IAA (A), PCIB (B), or NPA (C) was measured daily and plotted. Growth establishment of IR64 seedlings treated with IAA (D), PCIB (E), or NPA (F) was measured daily and plotted. *Error bars* indicate the S.E. of seedlings (n=10). Significance level was analyzed using One-way ANOVA; *P* < 0.01.

**Supplemental Figure S4.**
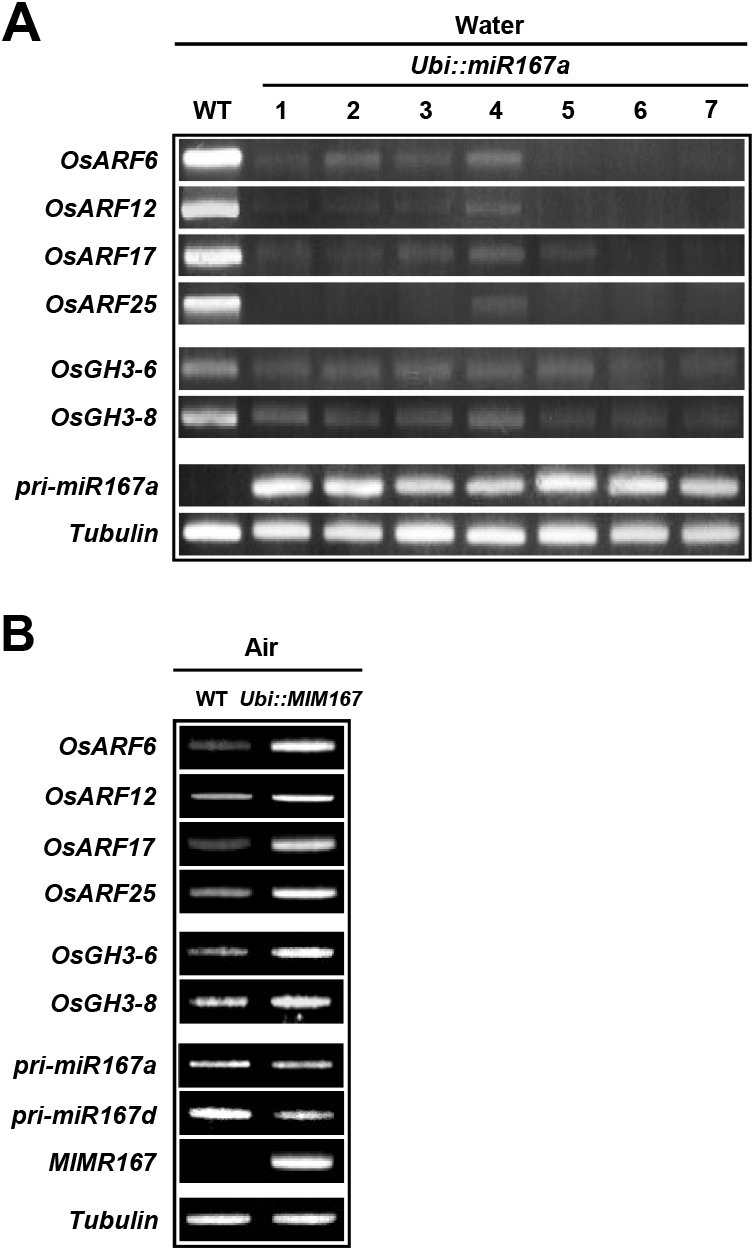
Expression of miR167 target genes and downstream-regulated genes in the wild type and transgenic rice coleoptile. Seeds from WT and T2 of homologous transgenic rice lines miR167a-1 to 7 carrying the *Ubi::miR167a* gene and miR167 target-mimic line MIM167 carrying the *Ubi::MIM167* gene transgenic rice were germinated and grown in the medium for 6 days under submerged (Water) or aerobic (Air) conditions. Total RNAs were purified from hypoxic and aerobic coleoptile and subjected to RT-PCR analysis. The expression of four target genes of miR167 (*OsARF6*, *OsARF12*, *OsAR17*, and *OsARF25*), two downstream-regulated genes (*OsGH3-6* and *OsGH3-8*), *pre-miR167a*, *pre-miR167d*, and *MIM167* in transgenic rice lines miR167a-1 to 7 (A) and miR167 target-mimic line MIM167 (B) and WT (A and B) were analyzed. The expression level of *tubulin* mRNA was measured and used as an internal control.

**Supplemental Figure S5.**
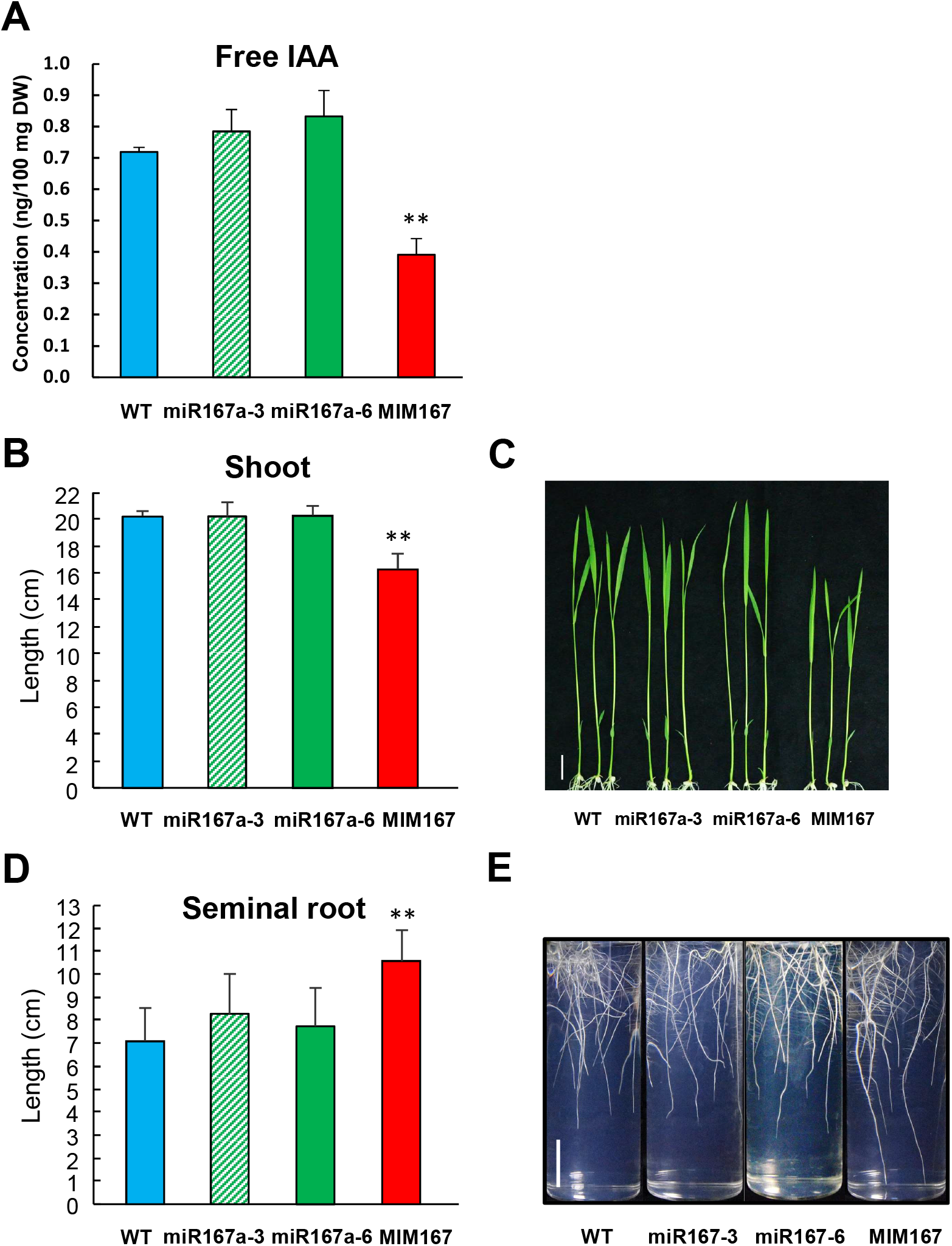
Effect of miR167 on the endogenous level of free IAA and the shoot and seminal root growth of aerobic seedlings. Three-day-old seedlings of wild type (WT) and transgenic lines miR167 (miR167a-3 and miR167a-6) and MIM167 carrying the *Ubi::miR167a* and *Ubi::MIM167*, respectively, were transferred to a medium solidified with 0.25% Phytagel and grown for 7 days at 28*°*C in a 16-h light (5,000 lux)/8-h dark cycle under aerobic conditions. Seedlings were then collected and the endogenous level of free IAA (A) was determined by UPLC-ESI-MS/MS analysis, and the length of the shoot (B) and seminal root (D) were determined using Image J software. Images of shoot (C) and root (E) of 10-day-old aerobic seedlings were shown. Scale bars: 2 cm. *Error bars* represent the S.E. of seedlings analyzed (n=20). Significance level between WT and transgenic lines was analyzed using Student’s *t*-test; “**” = *P* < 0.01.

**Supplemental Figure S6.**
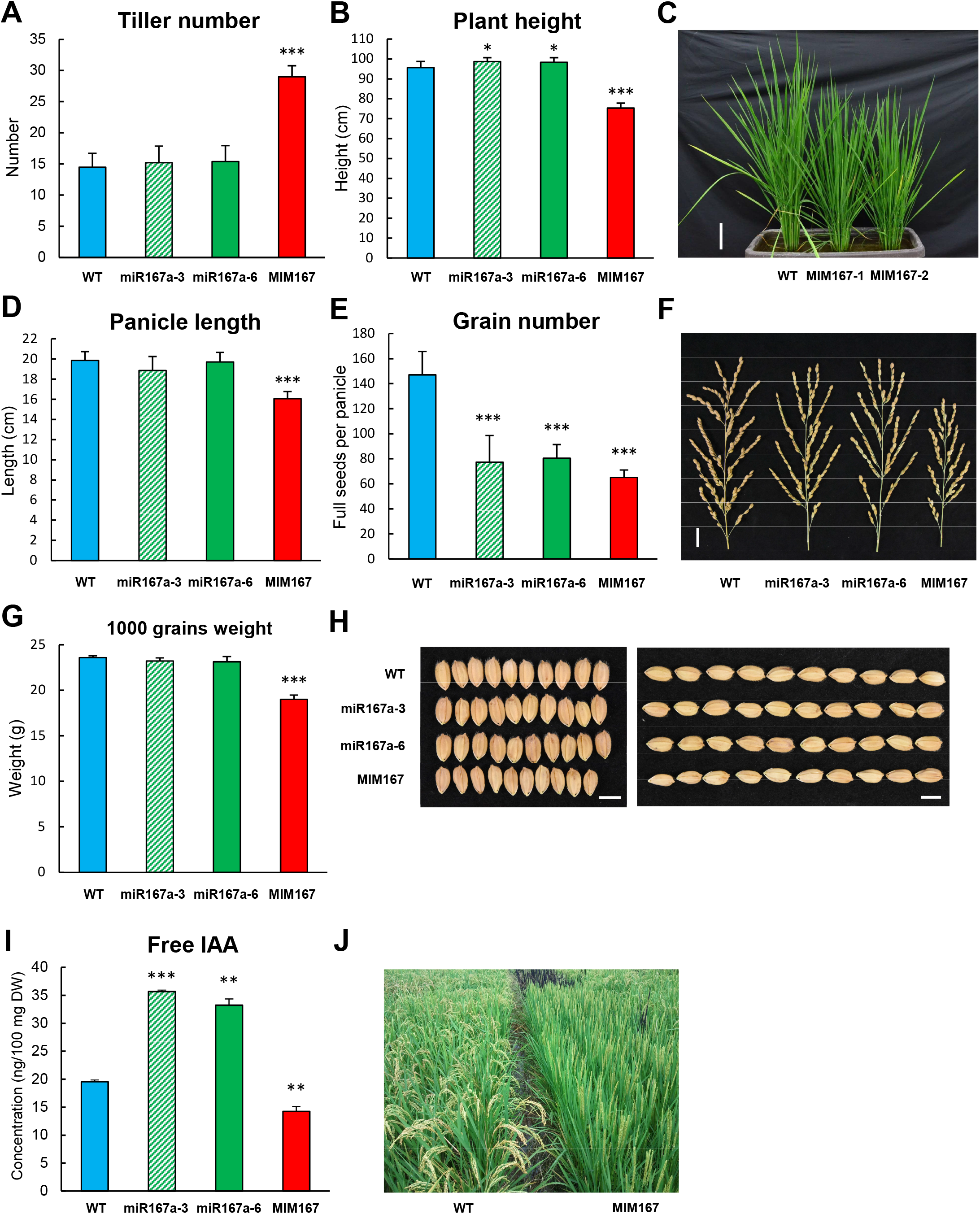
Effect of miR167 on the growth of mature rice plants. Seeds from wild type (WT), transgenic rice lines miR167 (miR167a-3 and miR167a-6), and MIM167 were germinated and grown in paddy fields for 150 days, except in (C). The number of tillers (A), plant height (B), panicle length (D), full seeds per panicle (E), and grains weight (G) were determined. (C) Image showing WT and MIM167 seeds germinated and grown in the greenhouse for 80 days. Scale bar: 10 cm. (F) Panicle structure of WT, miR167, and MIM167 lines. Scale bar: 2 cm. (H) Image showing the grain width (left of the panel) and length (right of the panel) of WT, miR167, and MIM167 lines (n=10). Scale bar: 5 mm. (I) Measurement of endogenous free IAA level from freshly harvested grains by UPLC-ESI-MS/MS analysis. (J) Plants morphology of WT and MIM167 grown in paddy fields. *Error bars* represent the S.E. (n=20). Significance level was determined using with the *t*-test; “**” = *P* < 0.01; “***” = *P* < 0.001.

**Supplemental Figure S7.**
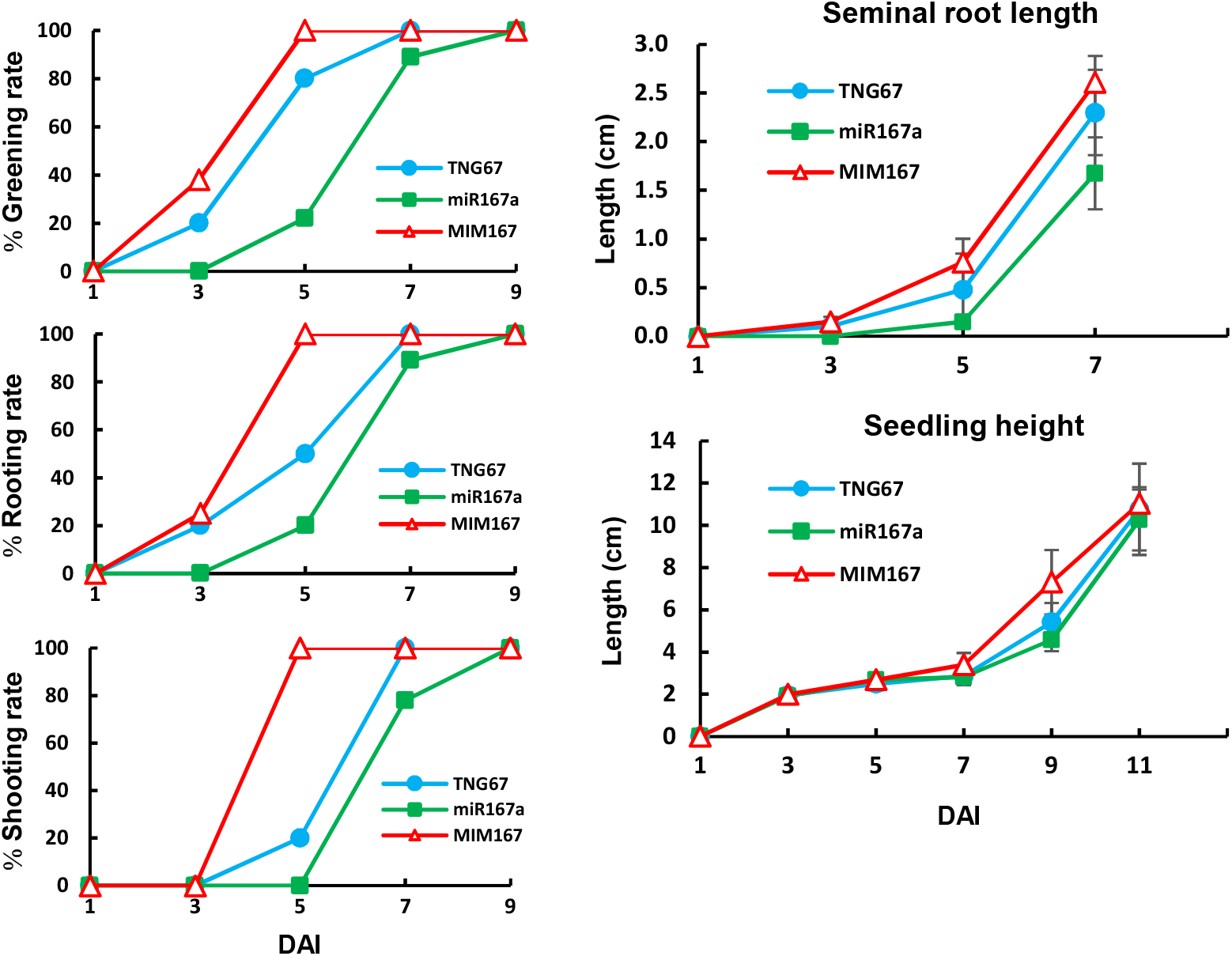
The target-mimic line MIIM167 exhibits enhanced tolerance to submergence than WT and miR167a overexpression line during AG. Seeds from WT, miR167a, and MIM167 were germinated and grown under fully embedded conditions, as described in Figure 7. The greening rate, rooting rate, shooting rate, seminal root length, and seedling height of hypoxic seedlings grown in sterile water containing 2.0 x 10^-2^ g/L phytagel at each time point were measured. *Error bars* indicate the S.E. of seedlings (n=10). Significance level was determined using One-way ANOVA, *P* < 0.01.

**Supplemental Figure S8.**
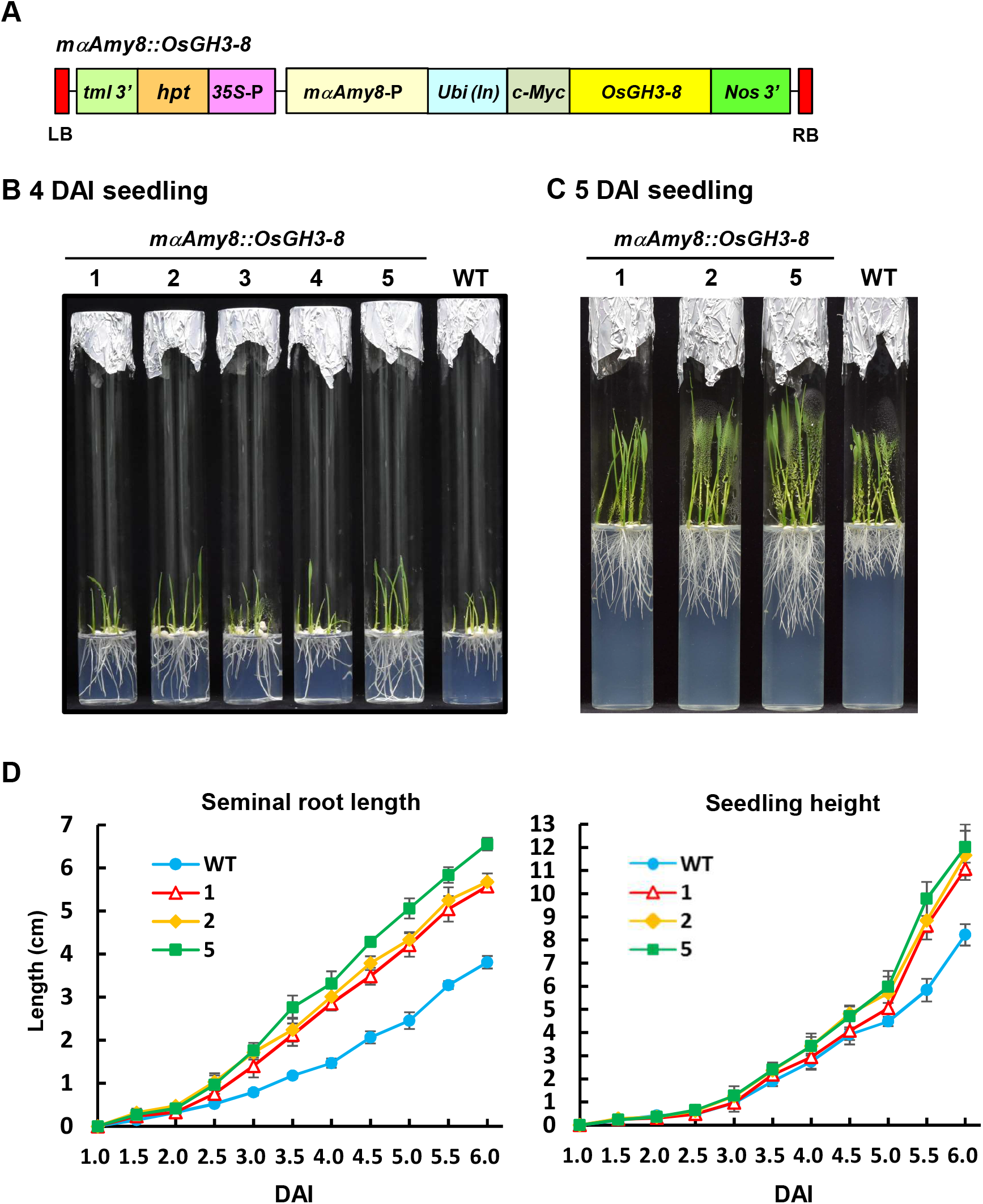
Overexpression of *OsGH3-8* leads to a significant increase in seminal root length of transgenic rice seedlings. (A) Schematic representation of construct of the binary vector used for rice transformation. The modified *αAmy8* promoter, *mαAmy8*, was used for the high-level expression of the *OsGH3-8* gene during seed germination. T2 seeds of homozygous transgenic rice lines 1-5 carrying the *mαAmy8::OsGH3-8* gene were obtained. Seeds from WT and transgenic rice were germinated and grown under aerobic conditions for 6 days. (B) Aerobic seedling morphology at 4 DAI and (C) 5 DAI. (D) The seminal root length of seedling and seedling height at each time point were measured. *Error bars* indicate the S.E. of seedlings (n=10). Significance level was determined using One-way ANOVA, *P* < 0.01.

**Supplemental Figure S9.**
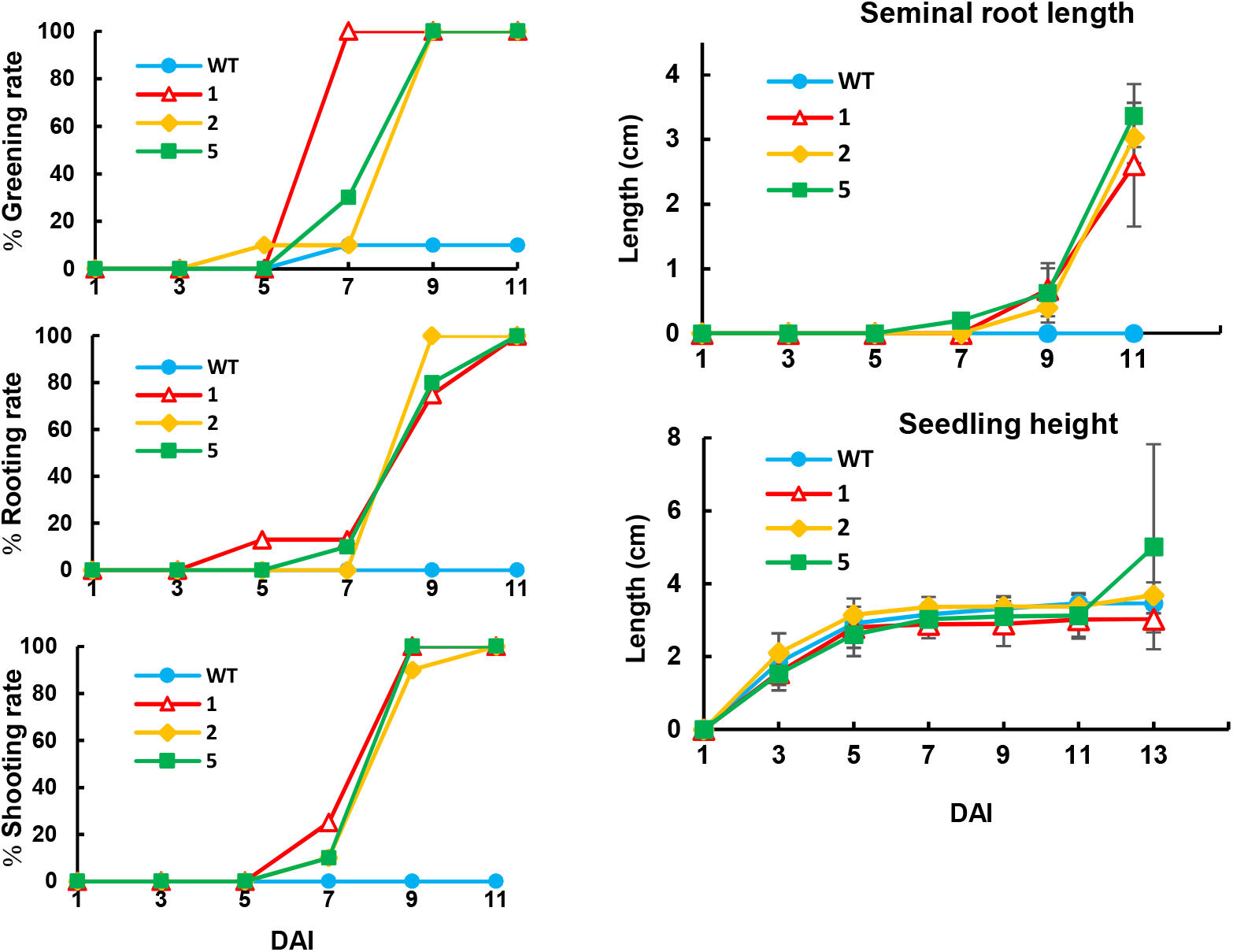
High-level expression of *OsGH3-8* in hypoxic seedlings confers enhanced tolerance to submergence during AG. T2 seeds of homozygous transgenic rice lines 1, 2, and 5 carrying the *mαAmy8::OsGH3-8* gene were obtained. Seeds from wild-type (WT) and transgenic rice were germinated and grown underwater, as described in Figure 7. Greening rate, rooting rate, shooting rate, seminal root length, and seedling height of seedlings grown in sterile water containing 3.0 x10^-2^ g/L phytagel at each time point were measured. *Error bars* indicate the S.E. of seedlings (n=10). Significance level was determined using One-way ANOVA, *P* < 0.01.

**Supplemental Figure S10.**
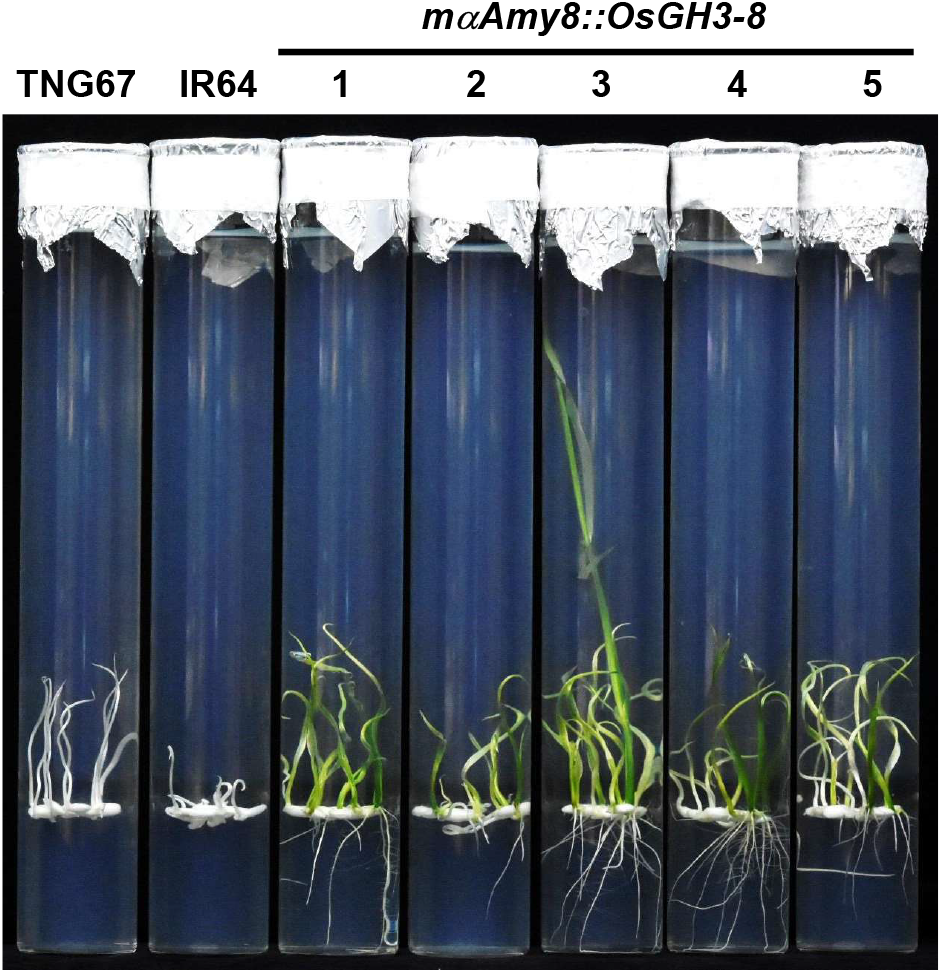
Overexpression of *OsGH3-8* under the control of the modified *αAmy8* promoter confers enhanced tolerance to submergence during AG in hypoxic seedlings. T2 seeds of homozygous transgenic rice lines 1 to 5 carrying the *mαAmy8::OsGH3-8* gene were obtained. Seeds from rice varieties TNG67, IR64, and transgenic rice were germinated and grown underwater, as described in Supplemental Figure S1. The test tube was filled with 70 mL of sterile water containing 4 mM CaCl_2_ and 2.5 g/L phytagel. Image of seedling morphology of TNG67, IR64, and *mαAmy8::OsGH3-8* at 15 DAI was shown.

**Supplemental Figure S11.**
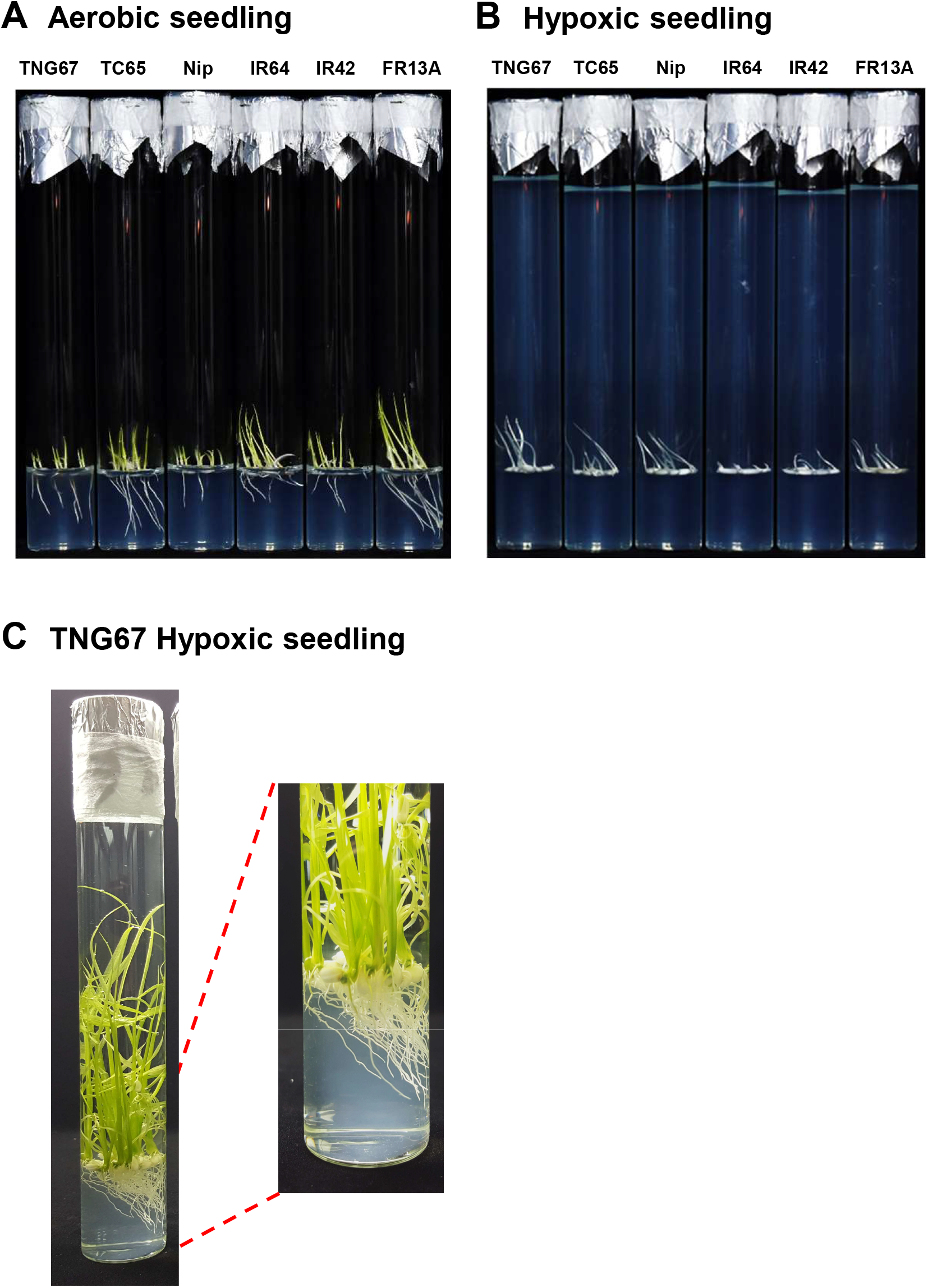
Rice coleoptiles are positively phototropic under submergence while roots are negatively phototropic. Seeds from three *japonica* varieties Tainung 67 (TNG67), Taichung 65 (TC65), and Nipponbare (Nip), and three *indica* varieties IR64, IR42, and FR13A were germinated and grown in light conditions (100 lux) in the air or under submergence for 3 days, and TNG67 hypoxic seedlings were grown for 28 days. Morphology of three-day-old Aerobic seedling (A) and hypoxic seedling (B) of six rice varieties was shown. Morphology of twenty-eight-old TNG67 Hypoxic seedling was shown in (C). Light source of low intensity was provided from the top left of the rice seedlings.

**Supplemental Table S1.**
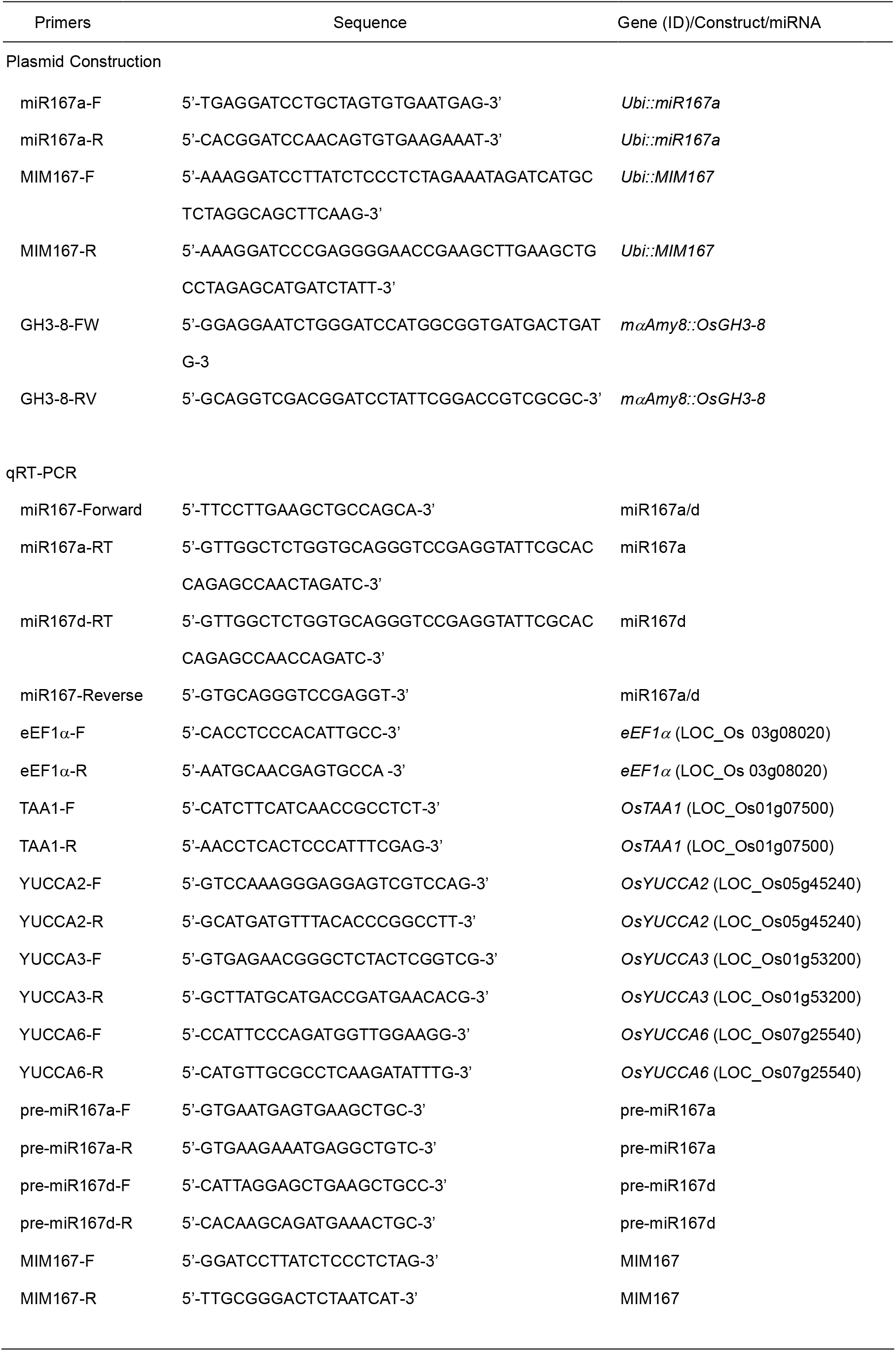
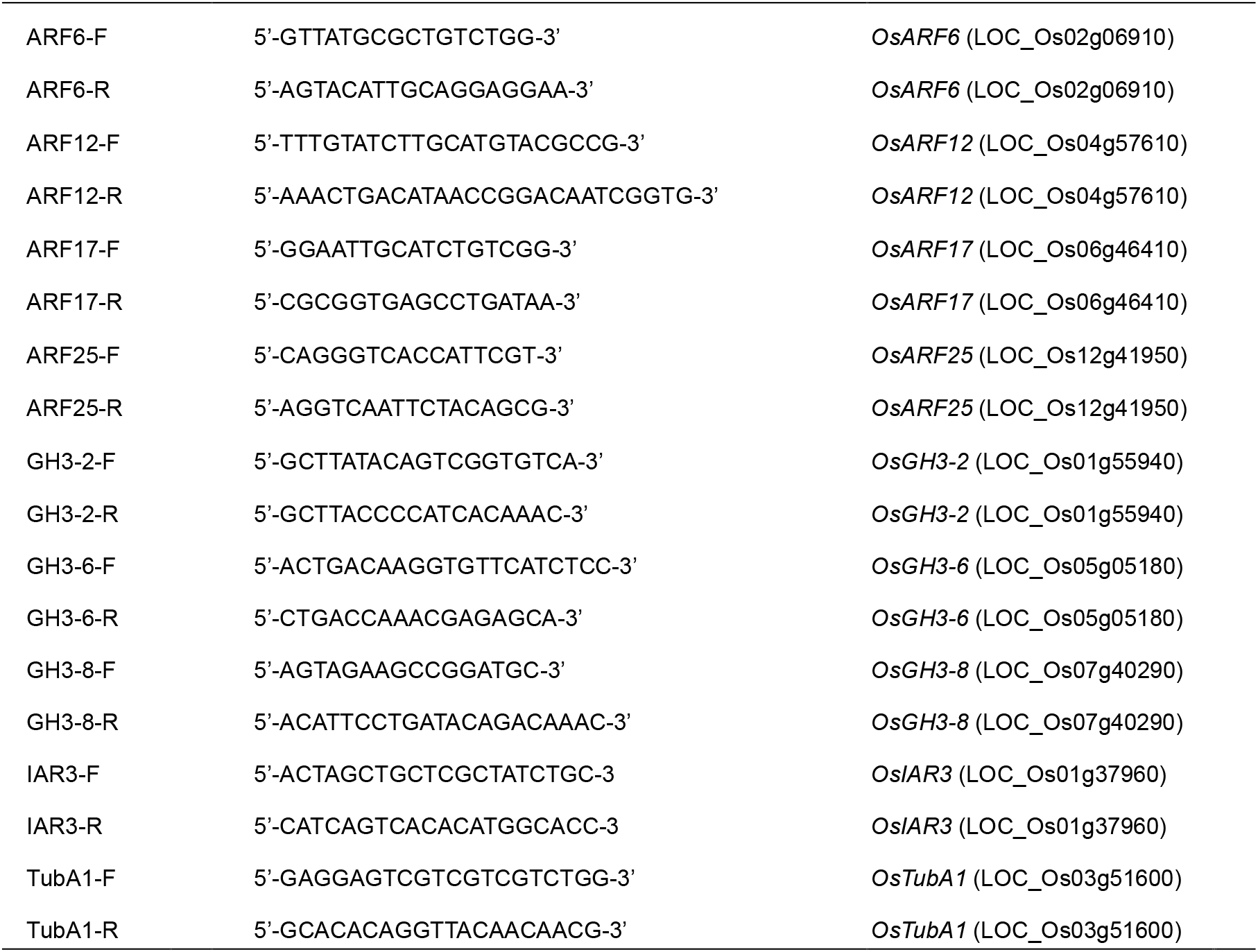
List of primers used in this study.

## Acknowledgments

We thank Dr. Carol Pei-Yin Wu for the critical review of the manuscript and Dr. Shih-Shun Lin’s technical assistance for small RNA gel blot analysis. We also thank Dr. Chih-Yu, Lin, and Ms. Ting-Hsiang, Chang for UPLC-ESI-MS/MS parameter optimization and Metabolomics Core Facility, Agricultural Biotechnology Research Center at Academia Sinica of Taiwan, for technical support.

## Author contributions

PWC conceived the project and designed the experiments; Jeremy JWC conducted the initial experiment to analyze miRNA microarray analysis and the stem-loop RT-qPCR experiments; KWL, CSW, HCC, HYC, HHK, YSL, YLC, HCC, SYS, and YCW performed the experiments; YCH advised on experiments; All authors analyzed the data. PWC and KWL supervised the conduct of the experiments and wrote the manuscript.

